# Nuclear BIN1 isoforms regulate c-Myc-mediated cell cycle control in oligodendrocytes

**DOI:** 10.1101/2025.09.30.676706

**Authors:** Iris Wai-Ting Ma, Gerald Wai-Yeung Cheng, Sunny Hoi-Sang Yeung, Martin Ho-Yin Yeung, Julia Kofler, Karl Herrup, Kai-Hei Tse

## Abstract

Bridging integrator 1 (BIN1) is a nucleocytoplasmic protein that inhibits c-Myc and acts as a tumour suppressor. BIN1 is ubiquitously expressed, but it is most abundant in skeletal myocytes and brain oligodendrocytes (OLs). BIN1 expression in the OL lineage is of particular interest, as the loss of myelin integrity is highly correlated with the progress of sporadic AD. More importantly, GWAS studies have identified rare BIN1 variants as the second strongest risk factor for sporadic Alzheimer’s disease (AD), after the ε4 variant of *APOE* gene. Despite these inherent interests, the control of the nucleocytoplasmic localisation as well as the modulation of the alternative splicing of the 20 exons of BIN1 are poorly understood in OLs. We report here the characterisation of BIN1 isoforms in OLs from two independent cohorts of postmortem AD brains using immunoblotting and immunohistochemistry and extend the findings to experimental APP/PS1 mice and primary murine OL cultures. Neuronal isoforms of BIN1 (BIN1:H, 95kDa) were significantly reduced (P < 0.0001), and the white matter/OL-specific isoforms (BIN1:L, 70kDa) were increased (P = 0.0349) in both AD cases and APP/PS1 mice. Importantly, the OL-specific BIN1 isoforms, identified by three different antibodies, were found in the nucleus of OL in human and mouse. Nuclear BIN1 was expressed by both the OL progenitor cells (OPCs) and mature OLs in vitro. Silencing Bin1 in OPCs led to a transcriptomic shift with a perturbed p53 pathway and cell cycle regulation, consistent with reduced Bin1-mediated c-Myc inhibition. The putative interacting sites between OL-specific BIN1:L and c-Myc were also identified by in silico analysis. The present findings suggest that nuclear BIN1 acts as a regulator of OL cell cycle control and support the hypothesis that BIN1 dysregulation in OL may contribute mechanistically to myelin pathology observed in sporadic AD.

## INTRODUCTION

Genome wide association studies (GWAS) identified Bridging Integrator 1 (*BIN1*) variants at the genetic loci conferring the second most significant risk factor for late-onset Alzheimer’s disease (AD) (Bellenguez et al., 2022; Chapuis et al., 2013; Harold et al., 2009; Seshadri et al., 2010). Only the APOE locus as a more powerful effect on AD risk. The major BIN1 variant allele (i.e. rs744373) is more frequently found in the general population (*BIN1* alleles frequency = 29% – 48.3%; Odd Ratio: 1.17 – 1.2) than the *APOE4* allele (Seshadri et al., 2010), but its expression pattern and function in the brain remain obscure. Encoded by 20 exons with a complex pattern of alternative splicing, BIN1 is a nucleocytoplasmic protein with twelve different isoforms. The nuclear functions of BIN1 were first identified in cancer cell where it serves as a MYC-inhibiting tumour suppressor (Elliott et al., 2000; Elliott et al., 1999). The cytoplasmic activities of BIN1 were later described in the brain (Tan et al., 2013). In AD patients, the BIN1 associated variants (e.g. rs744373; in the enhancer region) is associated with tau accumulation (Franzmeier et al., 2019), and the loss of cytoplasmic BIN1 function has been suggested to accelerate tau accumulation and propagation through impaired endocytosis (De Rossi et al., 2019; Ponnusamy et al., 2023; Taga et al., 2020; Zhang et al., 2024). Despite these findings, the cell type-specific expression, nucleocytoplasmic distribution and the differential functions of nuclear and cytoplasmic BIN1 in the brain are poorly understood, BIN1 is differentially expressed by the oligodendrocytes (OLs), astrocytes, neurons and their axons (Adams et al., 2016). Using well-defined monoclonal antibodies, strong nuclear BIN1 expression has been identified across different regions of the neuropil in human cerebral cortex and cerebellum (DuHadaway et al., 2003), where it is predominantly expressed by mature OLs, especially in the white matter tracts (De Rossi et al., 2016). However, these authors did not establish whether BIN1 is expressed in the OL nucleus, cytoplasm or both. In OL progenitor cells (OPC) and during their transition into myelinating mature OL, the transcription factor c-Myc controls cell cycle progression and regulates the epigenetic differentiation program (Jensen et al., 1998; Magri et al., 2014; Neumann et al., 2021). As Bin1 is a well-defined Myc inhibitor (Elliott et al., 1999; Pineda-Lucena et al., 2005; Sakamuro et al., 1996), here we hypothesised that BIN1 might be expressed in the OL nucleus where it could play a role in cell cycle regulation.

Here, using exon-specific primers and six well-defined commercial antibodies (De Rossi et al., 2016; DuHadaway et al., 2003; Wechsler-Reya et al., 1997), we examined the expression pattern of BIN1 transcript variants and protein isoforms in two cohorts of post-mortem human frontal cortices, in the cerebrum of transgenic AD mouse model APP/PS1 and in primary OL cell cultures derived from postnatal mouse. We confirmed the expression of nuclear BIN1 in the OL population and present evidence that Bin1 expression regulates the OL cell cycle. Together, the data show that BIN1, the second strongest genetic risk factor for sporadic AD, not only functions in neurons and tau processing (Dourlen et al., 2025; Zhang et al., 2024), but it also contributes to cell cycle regulation in the OL lineage.

## METHODS

### Human brain tissue study

Postmortem human brain tissue samples from the frontal cortex (Brodmann area 9) were obtained from two independent sources. The first cohort, comprising of frozen postmortem human brain tissues from the frontal cortex (Brodmann area 9) were obtained from the National Institutes of Health NeuroBioBank at the University of Maryland and the Human Brain and Spinal Fluid Resource Centre (Los Angeles, CA), following approval by the NeuroBioBank Tissue Access Committee. This cohort included a total of 24 cases comprising individuals diagnosed with non-Alzheimer’s disease (non-AD) dementias, Alzheimer’s disease (AD), and age-matched neurotypical controls (NCs). Cases with parkinsonian syndromes or major depressive disorder were excluded. All specimens consisted of unfixed tissue from the left frontal cortex, which had been snap-frozen and stored at –80°C until processing.

A second cohort of formalin-fixed, paraffin-embedded (FFPE) postmortem brain tissues was obtained from the Neuropathology Core of the Alzheimer’s Disease Research Centre at the University of Pittsburgh Medical Centre. Access to these tissues was granted under the oversight of the Committee for Oversight of Research and Clinical Training Involving Decedents at the University of Pittsburgh. This FFPE cohort also derived from Brodmann area 9 of the frontal cortex and included cases stratified by Braak neurofibrillary tangle stage, National Institute on Aging–Reagan Institute (NIA-RI) criteria, and APOE ε4 genotype. The cohort comprised neurotypical controls (NC: Braak stages 0–II; n = 9), sporadic AD cases (Braak stages III–VI; n = 17), and familial AD cases with confirmed *PSEN1* mutations (Braak stage IV; n = 7).

All protocols involving human brain tissue were approved by the Institutional Review Board of The Hong Kong Polytechnic University (HSEARS20200116004). Detailed demographic data, postmortem intervals, and additional case characteristics are provided in Supplementary Table S1.

### Animal Experimental Procedures

All experimental procedures involving animals were approved by the Animal Subjects Ethics Sub-Committee at The Hong Kong Polytechnic University (PolyU) (ASESC Case No.: 22-23/346-HTI-R-ECS-TSE) and conducted under a valid licence issued by the Department of Health, Hong Kong. Wild-type (WT) C57BL/6J mice and transgenic familial Alzheimer’s disease (fAD) model mice (B6;C3-Tg(APPswe,PSEN1dE9)85Dbo/Mmjax; MMRRC Strain #034829-JAX) were housed in individually ventilated cages (Techniplast, Italy) within a temperature-and humidity-controlled environment, maintained on a 12-hour light/dark cycle. Animals were provided with food (PicoLab Diet #5053, LabDiet Inc., MO, USA) and water ad libitum.

The APP/PS1 transgenic mouse model is a double transgenic strain that harbours a chimeric mouse/human amyloid precursor protein (*APP*) transgene with the Swedish mutation (APP_SWE_ K595N/M596L) and a human presenilin 1 (*PSEN1*) transgene with an exon-9 deletion (*PSEN1_dE9_*). This double-transgenic construct is integrated at a single locus on chromosome 9, regulated by the mouse prion protein (*Prnp*) promoter. The APP/PS1 model is well-characterised for developing widespread diffuse amyloid plaque deposits by 6 months of age (Jankowsky et al., 2001). Hemizygous APP/PS1 mice and age-matched wild-type controls were maintained until 18 months of age for experimental analyses.

At the experimental endpoint, animals were deeply anaesthetised and subjected to transcardial perfusion with cold phosphate-buffered saline (PBS), as previously described (Mok et al., 2023). Brains were rapidly dissected and bisected along the midline. The right hemisphere was fixed in 4% (v/v) paraformaldehyde for 24 hours, then cryoprotected in 30% (w/v) PBS-sucrose at 4°C for 72 hours. The fixed hemisphere was subsequently embedded and sectioned at 10 μm thickness using a cryostat, with sections collected onto charged slides for histological analyses. The left hemisphere was snap-frozen in dry ice and stored at –80LJ°C for subsequent analyses of gene and protein expression.

### Immunohistochemistry-Immunofluorescence

For human frozen tissue sections, slides were rehydrated by three successive washes in phosphate-buffered saline (PBS) and subsequently fixed in 4% (v/v) paraformaldehyde, as previously described. Fixed sections were blocked for non-specific binding using 10% (v/v) normal donkey serum in antibody diluent for 1 h at room temperature (RT). For murine tissue, frozen sections were washed and similarly blocked with 10% (v/v) normal donkey serum in PBS containing 0.3% Triton X-100 for 1 h at RT.

Blocked sections were incubated overnight at 4 °C with the following primary antibodies: anti-Olig2 (mouse monoclonal, 1:250, #MABN50, Millipore, CA, USA), anti-APC (Clone CC1, mouse monoclonal, 1:500, #MABC200, Millipore, CA, USA), and anti-BIN1 (rabbit polyclonal, 1:200, #ab185950, Abcam, Cambridge, UK). Following primary antibody incubation, sections were washed and then incubated with species-specific Alexa Fluor-conjugated secondary antibodies for 1 h at RT. Nuclear counterstaining was performed using 4’,6-diamidino-2-phenylindole (DAPI; Thermo Fisher Scientific, MA, USA), and slides were coverslipped with hydromount mounting medium (#HS-106, National Diagnostics, GA, USA). The full list of antibodies used in both immunohistochemistry and immunoblotting is provided in Supplementary Table S2.

Immunofluorescence imaging was performed at the University Research Facility in Life Sciences using a Nikon Eclipse Ti2-E Live-cell Fluorescence Imaging System equipped with a DS-Qi2 monochrome camera and a pE-4000 multi-wavelength LED illumination system (CoolLED, Andover, UK), incorporating blue (Ex: 380/55), green (Ex: 470/30), red (Ex: 557/35), and far-red (Ex: 620/60) excitation filters. Imaging was conducted using a Nikon Plan Apo λ 20× objective lens (numerical aperture 0.75; refractive index 1.0). For human sections, random fields (882.5 μm × 882.5 μm at 2424 × 2424 pixels) encompassing white matter (WM), grey matter (GM), and WM-GM boundaries were acquired. Whole-brain images were captured for murine tissue sections. All image data were processed and quantitatively analysed using QuPath v0.4.4 (Bankhead et al., 2017) and ImageJ (FIJI v1.53, NIH, MD, USA). Negative controls, in which the primary antibody was omitted, were included in all experimental runs.

### Immunohistochemistry-conventional

Conventional immunohistochemistry was performed on human formalin-fixed, paraffin-embedded (FFPE) brain sections using the BOND Rx fully automated staining platform (Leica Biosystems GmbH, Germany), as previously described (Cheng et al., 2022). Following deparaffinisation and rehydration, sections were incubated with three distinct anti-BIN1 antibodies: 99D (#sc-13575, mouse monoclonal, 1:750, Santa Cruz Biotechnology, CA, USA), 2F11 (#sc-23918, mouse monoclonal, 1:750, Santa Cruz Biotechnology, CA, USA), and ab185950 (rabbit polyclonal, 1:750, Abcam, Cambridge, UK). BIN1 antibody binding details is list in Figure S1. Immunoreactivity was visualised using the BOND Polymer Refine Detection Kit (Leica Biosystems, IL, USA), and nuclei were counterstained with Mayer’s haematoxylin in accordance with the manufacturer’s instructions. Omission of primary antibodies served as negative controls in all staining batches. Following staining, slides were dehydrated, coverslipped, and prepared for microscopic examination and digital image analysis. The whole-slide images (WSI) of immunostained sections were acquired using a NanoZoomer S210 digital slide scanner (Hamamatsu Photonics, Japan) at 40× magnification (resolution: 0.221 μm/pixel). Image analysis was performed using QuPath, as previously described (Cheng et al., 2023).

### Oligodendrocyte Cell culture

Primary oligodendrocyte (OL) cultures were established from postnatal day 2–6 (P2–P6) mouse pups, as previously described (Mok et al., 2023). Oligodendrocyte progenitor cells (OPCs) were cultured for 5 days and differentiated for 7 days. The murine OPC cell line Oli-Neu was also used (Jung et al., 1995). For loss-of-function studies, Bin1 expression in Oli-Neu cells was silenced using siGENOME SMARTpool Mouse Bin1 siRNA (Gene ID: 30948; #M-043588-01-0010, Horizon Discovery, USA) with DharmaFECT 1 Transfection Reagent (#T-2001-02, Horizon Discovery, USA). In the controls, cells were treated similarly using siGENOME SMARTpool Non-Targeting siRNA Pool #1 (#D-001206-13-05, Horizon Discovery, USA), as described (Yeung et al., 2024)

### Immunocytochemistry-Immunofluorescence

Immunocytochemistry was performed as described (Yeung et al., 2024). Primary OPC, OL or Oli-Neu cells plated on coverslips, were fixed with 4% (v/v) paraformaldehyde (pH 7.2) for 30 minutes, washed with phosphate-buffered saline (PBS), and blocked with 5% (v/v) normal donkey serum (#017-000-121, Jackson ImmunoResearch) in PBS with 0.3% (v/v) Triton X-100 for 30 minutes at RT. Cells were incubated for 2 hours with primary antibodies against BIN1, c-Myc or OL markers as listed in Table S2. Fluorescent visualisation was achieved using donkey anti-mouse or anti-rabbit Alexa Fluor-488,-555, or - 647 conjugated secondary antibodies (#A31571, #A31572, #A31570, #A31573, #A21206, #A21202, Thermo Fisher Scientific, MA, USA). Nuclei were counterstained with DAPI (1 µg/mL) for 5 minutes, and coverslips were mounted with hydromount. Imaging was performed using a Nikon Eclipse Ti2-E Live-cell Fluorescence Imaging System at the University Research Facility in Life Sciences. The system was equipped with a DS-Qi2 monochrome camera and a CoolLED pE-4000 multi-wavelength LED illumination system, with blue (Ex: 380/55), green (Ex: 470/30), red (Ex: 557/35), and far-red (Ex: 620/60) filters, using a Nikon Plan Apo λ 20× objective (N.A. 0.75). Duplicate coverslips were analysed per condition, with five random images (2424 × 2424 pixels) acquired from 882.5 µm × 882.5 µm fields, using Nikon NIS Elements software (V5.21.03). Image analysis was performed using QuPath.

### Western blotting

Cell or tissue lysates were prepared in radioimmunoprecipitation assay (RIPA) buffer (#20-188, Millipore, CA, USA) supplemented with cOmplete^TM^ protease inhibitor cocktail and PhosSTOP^TM^ (#11697498001, #4906845001, Roche, Hong Kong). Protein concentrations were quantified using the Bio-Rad Protein Assay (#5000001, Bio-Rad, CA, USA), normalised, and 30 µg of protein per sample was separated on 7.5–12% SDS-polyacrylamide gels. For denaturing conditions, samples were heated at 95°C for 5 min; for non-denaturing conditions, SDS and heat were omitted. Proteins were transferred to polyvinylidene difluoride (PVDF) membranes using the Trans-Blot Turbo system (#1704150, Bio-Rad, CA, USA) and blocked with 5% non-fat milk. Membranes were probed overnight at 4°C with primary antibodies BIN1 and related targets as listed in Table S2 Blots were incubated with horseradish peroxidase-conjugated secondary antibodies (Cell Signaling Technology), washed in Tris-buffered saline with Tween-20, and developed using SuperSignal™ chemiluminescent substrates (#34580, #34075, #34095, Pico, Dura, or Femto grade, Thermo Fisher Scientific, MA, USA). Signals were captured using a ChemiDoc MP Imaging System (#12003154, Bio-Rad, CA, USA) with 5-min exposures. Band intensities and molecular weights (kDa) were quantified using ImageJ (FIJI v1.53, NIH, MD, USA). Details of BIN1 binding sites are listed in Figure S1.

### Gene expression assay

Primers against human BIN1 variants and mouse Bin1 variants were designed using NCBI BLAST and the details are provided in Table S3. Other primer sequences were obtained from PrimerBank (Spandidos et al., 2010) or an earlier report (De Rossi et al., 2016). Total RNA was extracted with the RNeasy Mini Kit (#74106, Qiagen, Germany) and residual genomic DNA removed by DNase I digestion (1 U/µL, #18068-015, Invitrogen/Thermo Fisher Scientific, MA, USA). Reverse transcription was performed with the High-Capacity RNA-to-cDNA Kit (#4387406, Applied Biosystems, Thermo Fisher Scientific, MA, USA). Target genes and Gapdh were amplified by real-time PCR on a LightCycler 480 (Roche) using TB Green® Premix Ex Taq™ II (#RR820A, TaKaRa Bio Inc., Japan).

### RNA sequencing

For transcriptomic profiling of Bin1 knockdown in OPCs, total RNA (1000 ng) was processed with the Illumina Stranded mRNA Prep Ligation kit (#20040532, Illumina, CA, USA). Polyadenylated RNA was captured with RNA purification beads, fragmented with Elute, Prime, Fragment, High Concentration Mix (EPH3), and reverse transcribed using First Strand Synthesis Mix (FSA) followed by Second Strand Marking Master Mix (SMM; all from Illumina Stranded mRNA Prep Ligation kit, #20040532). After purification with AMPure XP beads (#A63881, Beckman Coulter Genomics, CA, USA), cDNA was adenylated at 3’ ends with A-Tailing Mix (ATL4) and ligated to Illumina RNA UD Index adapters (#20040553, Illumina, CA, USA) using Ligation Mix (#20040532, Illumina, CA, USA). Libraries were further purified with AMPure XP beads, amplified with Enhanced PCR Mix (EPM; #20040532, Illumina, CA, USA), and indexed with RNA UD Indexes Set A (#20040553, IDT for Illumina). Purified libraries were quantified with Agilent DNA HS chips (#5067-4626, Agilent Technologies, CA, USA) on a 2100 Bioanalyzer system. Sixteen libraries were pooled and sequenced on an Illumina NextSeq 2000 system with NextSeq 1000/2000 P2 reagent kit (#20040560; 200 cycles) in the University Research Facility in Life Sciences at PolyU. For RNA sequencing analysis, raw read files were aligned to mouse genome GRCm39 (Ensembl) using the Dragen-RNA application (v 4.3.4, BaseSpace). Aligned reads were subsequently processed for differentially expressed genes (DEGs) discovery using DESeq2 (v 1.42.1) within RStudio. The downstream investigation of the DEGs was performed using the Enrichr platform (Kuleshov et al., 2016) and the KEGG (Kanehisa et al., 2016), MSigDB (Molecular Signatures Database - Hallmark) (Castanza et al., 2023), Gene Ontology(Gene Ontology et al., 2023), PANTHER (Protein Analysis THrough Evolutionary Relationships) (Thomas et al., 2022), PPI-Transcription factor (Chen et al., 2013) and TRRUST v2 (Han et al., 2018) database therein. Protein-protein interactions were also analysed and graphed by STRING database (Szklarczyk et al., 2023).

### Structural analysis of BIN1 *in silico*

Sequences of BIN1 transcript variants and protein isoforms were retrieved from NCBI and Ensembl databases (Table 1). The experimentally validated BIN1 SH3 domain and c-Myc (Myc Box 1) interaction structure was obtained from RCSB Protein Data Bank (PDB: 1MV0; (Pineda-Lucena et al., 2005)). All other three-dimensional structures of human BIN1 and mouse Bin1 protein as well as c-Myc proteins were predicted by the Alphafold artificial intelligence algorithm deposited in the database (https://alphafold.ebi.ac.uk/; (Jumper et al., 2021)), as listed in Table S6. To evaluate the potential interactions between the whole protein structures of human BIN1 and c-Myc, as well as murine Bin1 and c-myc, these models were systematically docked by using ClusPro web server 2.0 (Kozakov et al., 2017). The computed binding affinity between the protein structures were ranked based on scores measured by the estimated free energy coefficient (*E_hydrophobic_*) and the binding cluster size at the centre. The best ranked protein-protein interaction structures were then explored, with molecular graphics, annotation and analyses performed by using software ICM Browser Pro (Molsoft L.L.C. (Abagyan et al., 1994)) and UCSF ChimeraX (Meng et al., 2023).

**Table 1.**
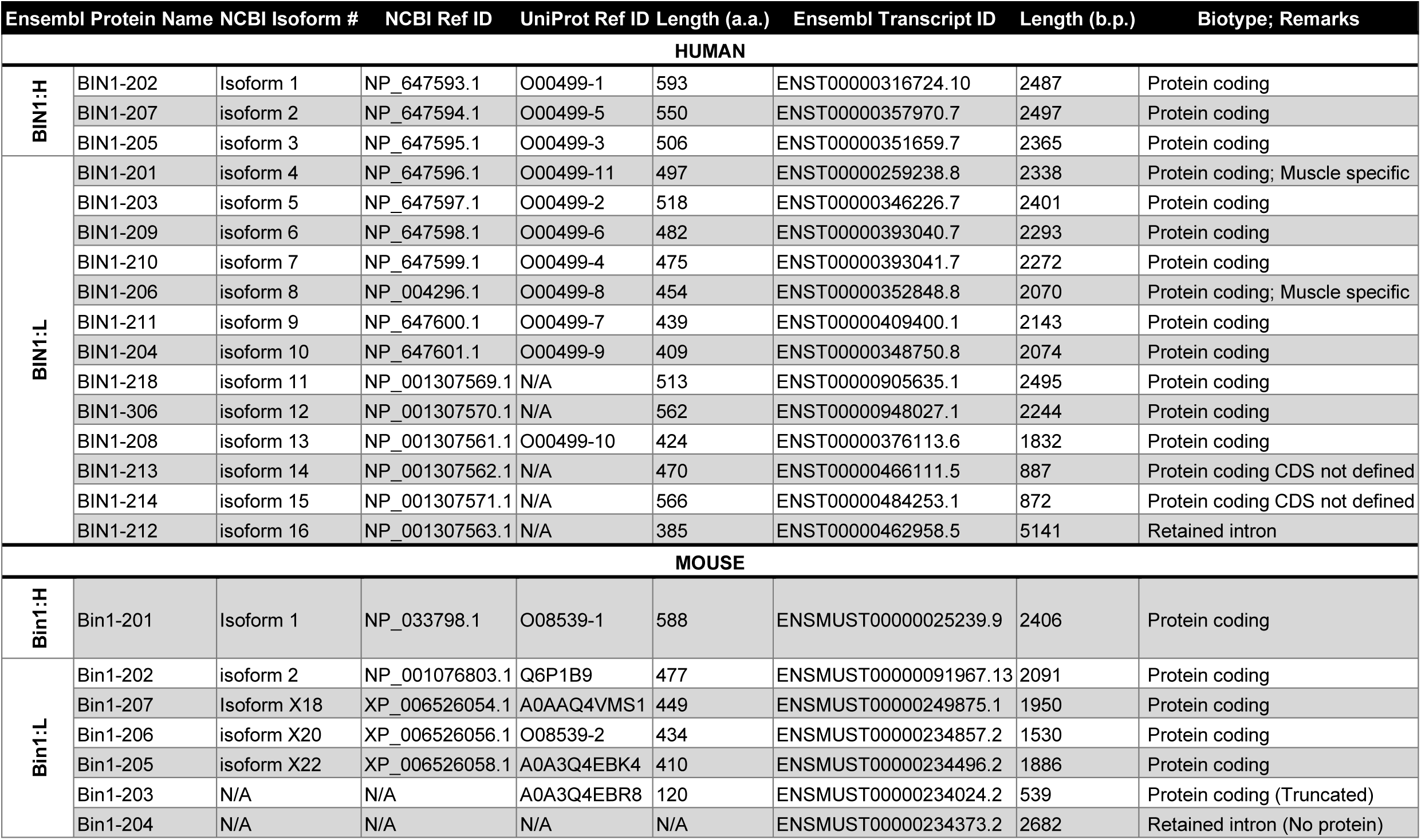
Human and mouse BIN1 isoforms and nomenclature.

### Statistical Analysis

Experiments comprised a minimum of three independent biological replicates, with the number of experiments specified in each figure. All results presented as means ± standard error of the mean (SEM). Outliers were identified using Grubbs’ test. Comparisons between two groups were performed with unpaired t-tests; one-way ANOVA with post hoc multiple comparisons was applied for single-variable analyses across more than groups. For analyses involving two or more variables across multiple groups, two-way ANOVA with post hoc testing was used. Statistical analyses were performed in GraphPad Prism v10.6 (GraphPad Software Inc., MA, USA) and details provided in figure legend.

## RESULTS

### Characterisation of BIN1 isoforms homology and antibody targets

The human BIN1 protein is encoded by 20 *BIN1* exons whose inclusion in the final mRNA transcript is regulated by alternative splicing (De Rossi et al., 2016; Taga et al., 2020; Tan et al., 2013) resulting in thirteen possible protein isoforms (Fig. 1A, Table 1). BIN1 proteins are conserved among mammalian species with a high homology (95.74%, isoform 1) between human (593 a.a.) and mouse (588 a.a.). In both human and mouse, BIN1 protein isoforms share a general structure characterised by a BAR domain at the N-terminus (Exon 1 - 10) of the protein, followed by a PI domain (phosphoinositide binding module; Exon 11), a CLAP domain (clathrin and AP2 binding; Exon 13 - 16), the MBD domain (myc-binding domain; Exon 17, 18), and a SH3 domain at the C-terminus (src homology 3; Exon 19, 20) (Fig. 1A). The differential splicing of Exon 7 in the BAR domain results in the subdivision of BIN1 protein isoforms into two groups: BIN1:H (**<u>H</u>**igh molecular weight; isoforms 1-3 or Ex7) and BIN1:L (**<u>L</u>**ow molecular weight; isoforms 4-12; or D7), respectively (De Rossi et al., 2016). In the human brain, BIN1:H isoforms are found in neurons and astrocytes (Taga et al., 2020; Toga et al., 2015), while BIN1:L isoforms are primarily associated with OLs especially in the WM (De Rossi et al., 2016). BIN1 isoform 1-3, 5-7, 9, 10 and 13 are expressed in human brain (Butler et al., 1997; Taga et al., 2020; Tan et al., 2013). In mouse, Bin1:H (isoform 1) and Bin1:L (isoforms 2, X20, X22) can be found in the brain. A comprehensive table summarising the isoforms and transcript variants can be found in Table 1. To investigate the expression pattern and function of BIN1 isoforms in OLs, we first fully characterised the binding sites of six commercially available antibodies and predicted their putative BIN1 isoform detection in human and mouse (Fig. 1A and Fig. S1). Briefly, these antibodies are directed at epitopes in the BIN1 at the BAR (2F11, CST13679), MBD (99D, G10) and SH3 (ab185950, 1H1) domains and detect different BIN1 isoforms, respectively. Note in particular that antibody clones 99D and 2F11 detects and distinguish isoforms with (BIN1:H) or without (BIN1:L) Exon 7 respectively, in both human and mouse (Fig. S1) (DuHadaway et al., 2003; Wechsler-Reya et al., 1997).

**Figure 1.**
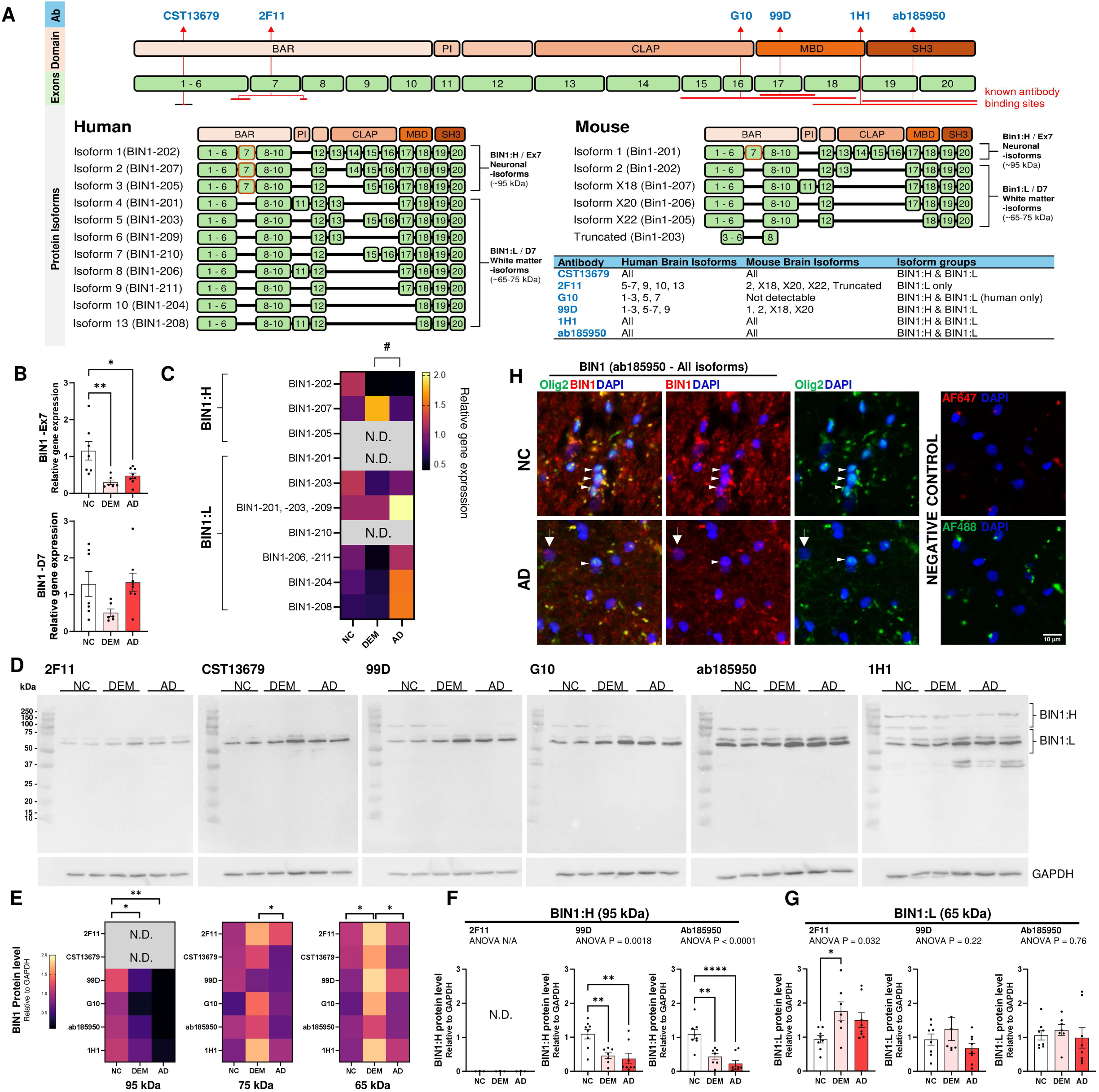
BIN1 isoforms structures and their differential expression in Alzheimer’s disease brain tissues. **A** The schematic structure of human and mouse BIN1. *Above*: the protein functional domains (orange boxes) of BIN and corresponding exon number (green boxes): BAR (membrane binding), PI (phosphoinositide binding); CLAP (AP2 and clathrin binding in endocytosis); MBD (Myc-binding domain) and SH3 (interact with proline rich proteins including c-Myc). *Below left* Structures of the 11 common human BIN1 isoforms (BIN1:H: Isoform 1-3 with exon 7, neuronal-specific; BIN1:L Isoforms 4-13 without exon 7, glia-enriched). *Below left* Structures of the 6 common mouse BIN1 isoforms. Target epitopes of the six BIN1 antibodies used in this study are pointed by arrows with targeted isoforms in human brain summarised in the blue box and binding site sequence listed in Fig. S1 **B** Gene expression analysis with published primers (De Rossi et al., 2016) showed a significant reduction in BIN1 transcripts containing exon 7, but not without, in DEM and AD as compared with NC (One-way ANOVA, Tukey post-hoc, *P < 0.05; **P < 0.01) in frozen human brain tissues. **C** BIN1 variant-specific primers identified a significant and divergent difference across NC, DEM and AD groups (P = 0.0281) and transcripts (P = 0.0376), with a significant pairwise difference between DEM and AD (Two-way ANOVA, Šídák’s post-hoc, ^#^P < 0.05). **D** Representative western blots of BIN1 expression using antibody clones of 2F11, CST13679, 99D, G10, ab185950 and 1H1 which showed a consistent downregulation of BIN1:H (≥95 kDa) and an upregulation of BIN1:L (≤65 kDa) in DEM and AD groups. GAPDH served as loading control. **E** Heatmaps summarising the quantifications of BIN1 isoforms at 95 kDa, 75 kDa, and 65 kDa confirming the observations (Two-way ANOVA, Šídák’s post-hoc, *P < 0.05; **P < 0.01) **F, G** Bar charts of BIN1:H (95 kDa) and BIN1: L (65 kDa) expression detected by 2F11, 99D and ab185950 confirming their significant and divergent regulation in DEM and AD groups NC (One-way ANOVA, Tukey post-hoc, *P < 0.05; **P < 0.01; ****P < 0.0001; Overall ANOVA P values as denoted). N.D.: Not detectable. **H** Representative immunohistochemistry-immunofluorescence images showing the presence of BIN1-IR (ab185950; red) in the Olig2^+^ OL cells (arrowheads; green) and Olig2 negative cell (arrow) in NC and AD. Negative control with antibodies omitted as shown.

### BIN1 expression in Alzheimer’s disease brain

We used specific PCR primers to investigate the differential transcription of *BIN1* isoforms in the frontal cortex of a cohort of non-AD dementia (DEM), AD or age-matched control cases (Table S1). Using specific primer pairs from an earlier study (De Rossi et al., 2016), we found that BIN1:H isoforms (Exon 7 (Ex7)-specific) were significantly reduced in all dementia cases, but BIN1:L isoforms (Exon 7-skipped (D7)) were differentially regulated in DEM and AD samples (Fig. 1B). By designing primers specifically targeting the boundary of the alternately spliced regions across BIN1:H and BIN1:L transcripts (see Table S3), we confirmed the differential expression (Fig. 1C; P = 0.0281), and the consistent absence (N.D.) of isoform 3 (BIN1-205) and 4 (BIN1-201) in human brain (Taga et al., 2020).

These divergent *BIN1* expression pattern were reflected at the protein level, as characterised in western blots using the six different antibodies (2F11, CST13679, 99D, G10, ab185950 and 1H1; Fig. 1D, Fig. S1). Based on the molecular weight, the BIN1:H isoforms (95 kDa) detected with 99D, G10, ab185950 and 1H1, were significantly reduced in both DEM and AD subjects (Fig. 1E). In contrast, BIN1:L isoforms (65-75 kDa) predominated among all cases and were upregulated. BIN1:L is a WM-specific group of BIN1 isoforms (De Rossi et al., 2016). Quantifications of the immunoblots confirmed that 2F11 was specific for human BIN1:L isoforms and was increased by disease conditions (Figure 1F, G). To visualise the cellular localisation of BIN1, immunohistochemistry was performed. The antibody clone ab185950, a pan-BIN1 antibody, detected BIN1 immunoreactivity colocalising with Olig2^+^ cells in both control and AD cases (Fig. 1H). These observations were consistent with the single-cell RNA-seq databases of human and mouse brains, where *BIN1* expression was most significantly enriched in the mature OL population (Table S4) (Karlsson et al., 2021; Saunders et al., 2018; Uhlen et al., 2015).

### BIN1 is expressed in the nucleus and cytoplasm of oligodendrocytes in Alzheimer’s disease

We next examined BIN1 nuclear localisation pattern in the frontal cortex of a second cohort of human cases with known sporadic AD (sAD), hereditary familial AD (fAD) and age-matched controls, which were characterised by Braak stages using antibodies clones ab185950, 99D and 2F11 antibodies (Fig. 2A-F, and Table S1). At low magnification, all three antibodies revealed a strong BIN1 presence in the WM with BIN1^+^ bundles that bore a high resemblance to cortical myelinated fibres coursing into GM (Fig. 2A-C). At high power, cytoplasmic BIN1-IR presented in the neuropil as bundles of fibres. In addition, we observed cells resembling glia with clear nuclear BIN1^+^ staining throughout the cortical layers (Fig. 2D). In the GM, cells with BIN1^+^ nuclei were usually small with highly condensed chromatin and positioned adjacent to cortical neurons. Thus, these cells strongly resemble perineuronal OLs. A few cells with larger BIN1^+^ nuclei with less condense chromatin were found, presumably representing astrocytes were also observed (Fig. 2D). The nuclei of cortical neurons across all layers were negative for BIN1-IR. In the WM, apart from the heavy BIN1 labelling of the myelin fibres, columns of cells resembling OL with condensed BIN1^+^ nuclei were frequently observed.

**Figure 2.**
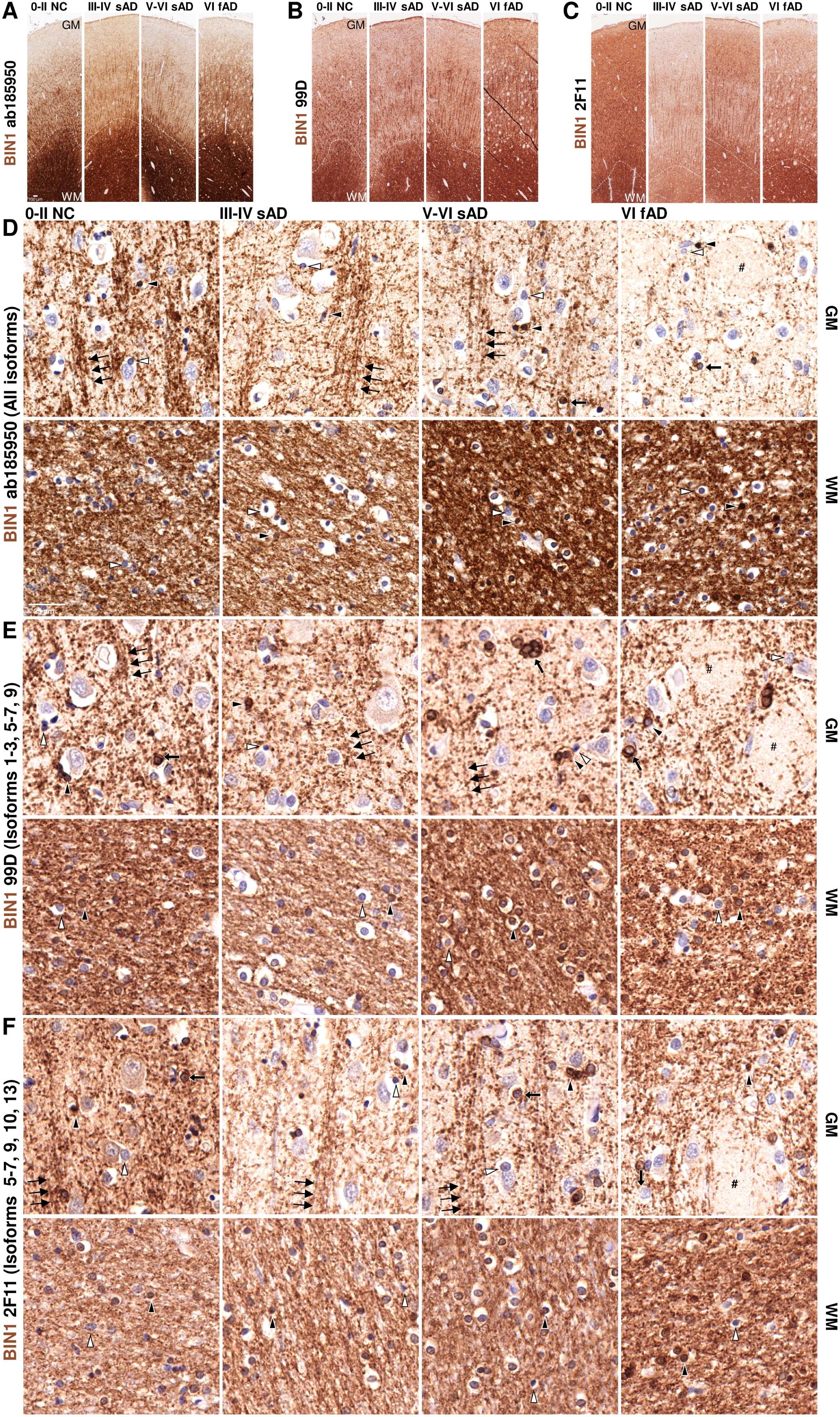
Nucleocytoplasmic BIN1 expression pattern in human AD brains detected by three distinct antibodies. **A-C** Representative conventional immunohistochemistry images across the frontal cortices of NC and AD cases with different Braak stages, including 0-II NC, III-IV AD, V-VI AD, and VI familial AD using antibody clones **A** ab185950 (targeting SH3 domain), **B** 99D (targeting the MBD domain), and **C** 2F11 (targeting the exon 6-exon junction at the BAR domain) at low power (DAB). All antibodies similarly demonstrated a pattern in the grey matter (GM) and white matter (WM), which was more prominent in the latter (GM-WM boundary indicated by dash lines). **D-E** Representative conventional immunohistochemistry images of GM and WM, with the use of antibody clones **D** ab185950, **E** 99D, and **F** 2F11 shown in high power. All antibodies showed a similar pattern of BIN1 expression, which was primarily found along bundles of myelinated axon-like fibres in the GM (triple black arrows). In both the GM, BIN1 expression was also found in the nucleus of OLs (black arrowheads) that sits adjacent to large neuronal cell body as satellite cells, as well as in the surrounding neuropil. These nuclear BIN1^+^ OLs were found along the threads of multiple aligned small nuclei bundles (black arrowheads) in the WM. BIN1 negative OL nuclear are indicated for comparison (white arrowheads). In both GM and WM, nuclear BIN1 was also found in heterochromatic nuclei resembling astrocyte-like cells (thick black arrows). In AD cases with advanced Braak stage (V-VI), the BIN1 immunoreactivity was minimal in the empty regions resembling prominent protein aggregations as denoted by # in **D, E, F**, which can be observed at low power (**A-C**, VI fAD). In all cases and antibodies, no neuronal BIN1 expression was found.

In advanced AD cases, BIN1 immunoreactivity was reduced, and the decrease was observed with all three antibodies in the plaque occupied area of both sAD and fAD, in agreement with an earlier report (Adams et al., 2016). The nucleocytoplasmic locations of the different BIN1 isoforms were similar in both GM and WM and the pattern was reproduceable across all three BIN1:L antibodies (Fig. 2D-F, quantifications in Table 2). In the GM, a significant reduction of cytoplasmic BIN1-IR was found in both sAD (Braak III-IV and Braak V-VI) and fAD (Braak VI) cases, a finding confirmed with all three antibodies across neuropil (P = 0.0109; Table 2). The density of cells with BIN1^+^ nuclei was also decreased and was sensitive to the increasing Braak stage (P = 0.0163; Table 2). No changes in BIN1-IR and or cells with BIN1^+^ nuclei were found in the WM. A Pearson coefficient test demonstrated a significant correlation between the nuclear BIN1^+^ cell density detected by the three BIN1 antibodies in the GM, but not in the WM (Fig. 3). As the three BIN1 antibodies target different isoforms (Fig. 1), this finding suggests that WM BIN1 isoforms maybe more diverse than those found in the GM.

**Figure 3.**
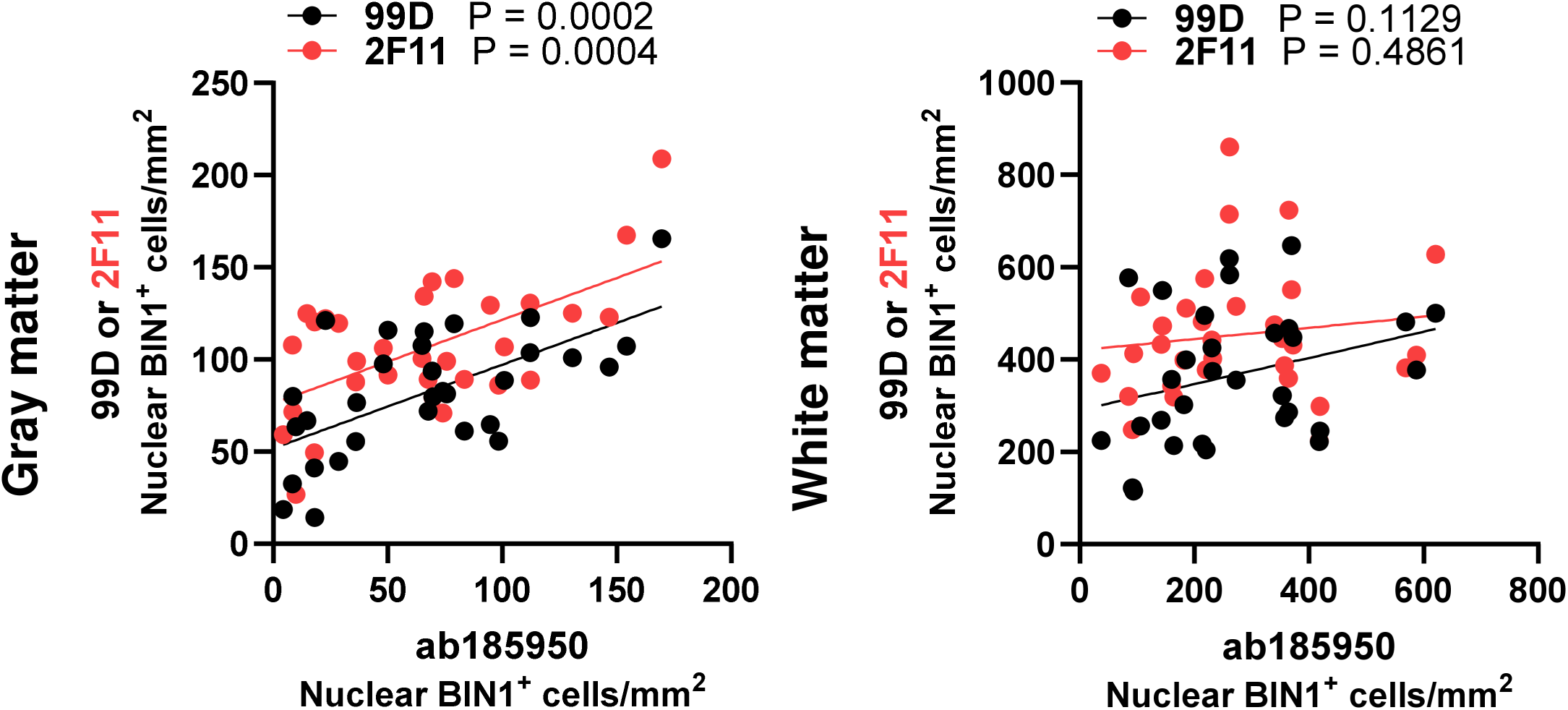
Correlation of BIN1 isoform protein expression in human frontal cortex. Correlation tests of the density of BIN1-positive cells between 99D or 2F11 and Ab185950 in GM and WM. Pearson test for correlation, P value as indicated.

**Table 2.**
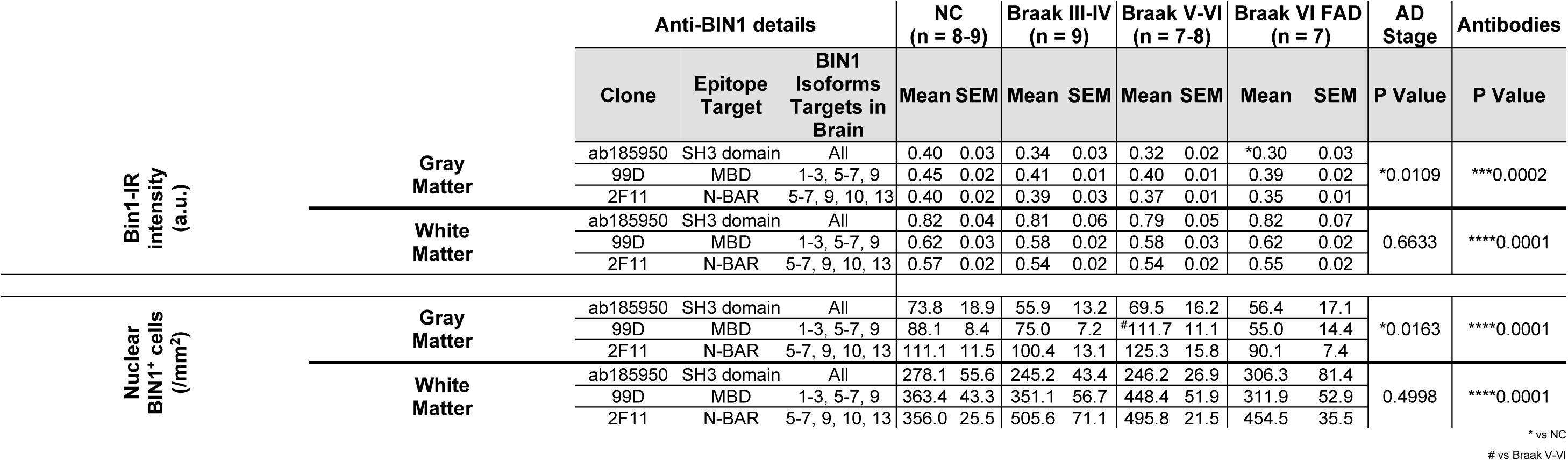
Quantification of BIN1 immunohistochemistry in human Quantifications of BIN1-immunoreactivity(-IR) and BIN1-positive nuclei across the frontal cortices of NC and AD cases with different Braak stages, including 0-II NC, III-IV AD, V-VI AD, and VI familial AD. Two-way ANOVA with Šidák post-hoc test (overall P value as annotated; Pairwise comparison vs NC, *P < 0.05; vs Braak VI FAD, ^#^P < 0.05).

### Cytoplasmic and nuclear Bin1 isoforms expression in oligodendrocytes in the mouse brain

There is a substantial structural homology between the amnio acid sequences of human and mouse BIN1. Not surprisingly, therefore, when we mapped the binding sites of the six antibodies, we found a high degree of epitope conservation, except for the G10 antibody (Fig. 1A, Fig. 1S). While cytoplasmic Bin1 localisation was reported in mouse brain previously (Ponnusamy et al., 2023), immunostaining using the pan-Bin1 antibody, ab185950, we found a significant presence of Bin1 in the nuclei of small cells resembling OLs across different brain regions (Fig. 4A, B). In GM, these small cells were juxtaposed to neurons (Fig. 4A); in WM tracts, nuclear Bin1^+^ cells were arranged in columns or threads across the corpus callosum, anterior commissure or pyramid tract near spinal cord (Fig. 4B). All these Bin1^+^ cells bore strong morphological resemblance to OL. Bin1 was commonly found in both nucleus and cytoplasm cells in all regions, but nuclear expression was nearly absent in hippocampus (Fig. 4A). Immunohistochemistry-immunofluorescence using antibodies against Olig2 confirmed that the Bin1^+^ cells belonged to the OL lineage (Fig. 4C, D), while CC1 immunostaining demonstrated that the majority of Bin1^+^ cells in both GM and WM were mature OLs (Fig. 4D, E). NeuN^+^ neuronal nuclei were negative for Bin1, as in human brain.

**Figure 4.**
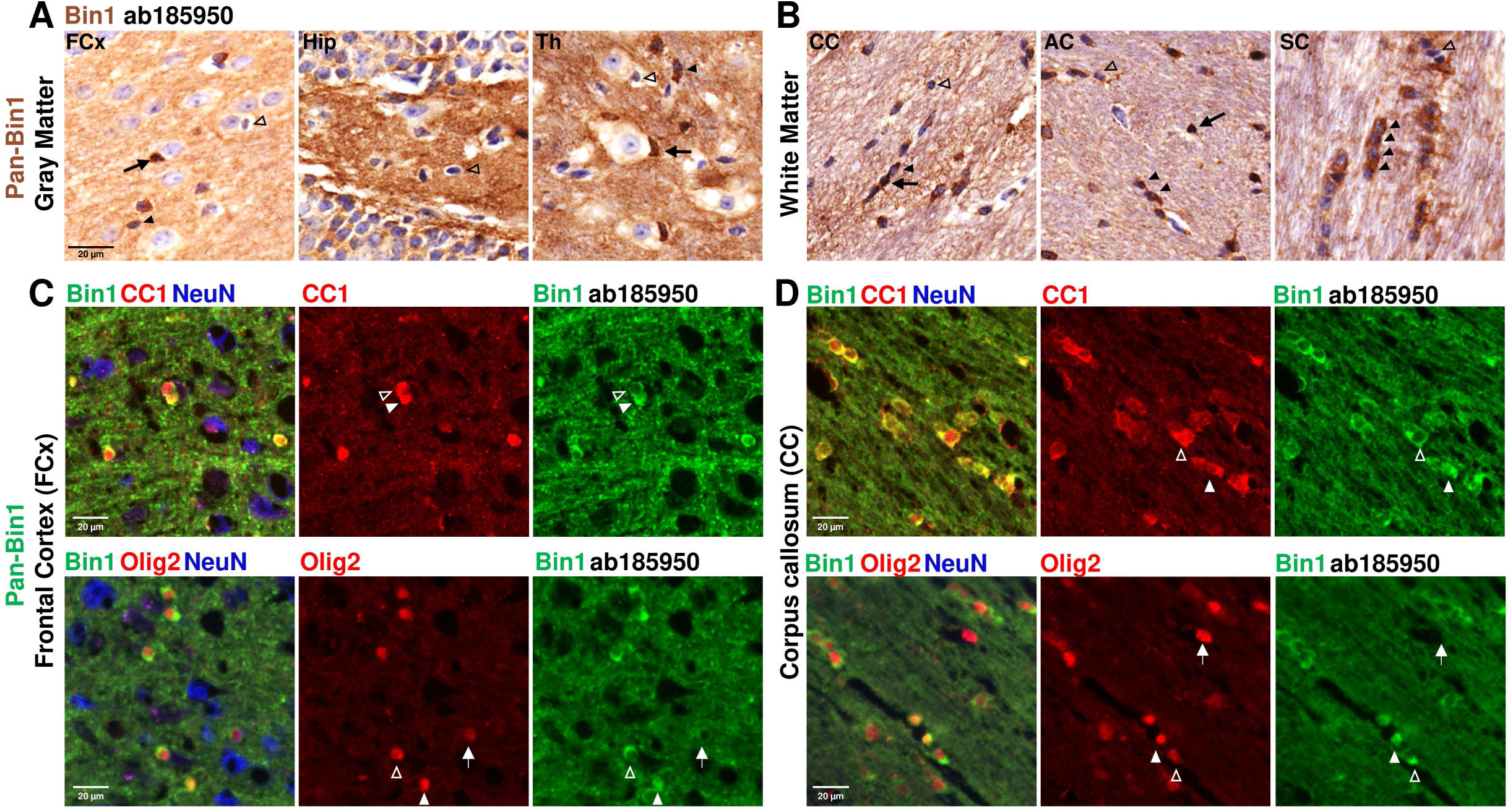
Nuclear BIN1 protein expression in mature oligodendrocytes in mouse tissue. Representative conventional immunohistochemistry images of pan-Bin1 antibody (ab185950; DAB) across **A** GM including FCx (frontal cortex), Hip (hippocampus), and Th (Thalamus), and across WM including **B** CC (corpus callosum), AC (anterior commissure), and SC (spinal cord) at Bregma 0.8 in WT mice brain tissues. In GM, nuclear Bin1^+^ OL was identified as the condensed nuclei adjacent to Bin1 negative neurons (black arrows). Cytoplasmic Bin1^+^ OL was also observed (black arrowheads), as compared to Bin1 negative OLs (open arrowheads). In WM, nuclear and cytoplasmic Bin1^+^ OL were similarly identified, where multiple of BIN1^+^ cells were found as a thread in myelinated tracts (quadruple black arrowheads in SC). **C, D** Representative immunohistochemistry-immunofluorescence images demonstrating the colocalization of *Above* Bin1 (ab185950; green) with NeuN (blue), and CC1 (red), or *Below* of Bin1 (ab185950; green), NeuN (blue), and Olig2 (red) in the brain of in **C** FCx and **D** CC in WT mice. In both regions, cytoplasmic (open arrowheads) and nuclear Bin1 signal (white arrowheads) was found in both the CC1^+^ mature OL and the Olig2^+^ OL lineage. Bin1 negative Olig2^+^ OL was also identified (white arrows). No NeuN positive neuronal nuclei were positive for Bin1-IR.

To further investigate the relationship of BIN1 isoforms and amyloid deposition, we examined brain tissues from APP/PS1 transgenic mice, a commonly used model of familial AD. They harbour in their genome both the transgenic construct of human *APP* cDNA with the Swedish mutation (APP^Swe^) as well as the human *PSEN1* with the deletion of exon 9 (PS1^ΔE9^). Using the 6E10 antibody, we observed heavy amyloid plaque deposition in the aged APP/PS1 mice across the cortical and subcortical regions (Fig. 5A). In both the APP/PS1 and wild-type mice at 18-month-old, cytoplasmic Bin1^+^ OLs were identified across cerebral cortex and corpus callosum (Fig. 5B, C, open arrowheads). In the APP/PS1, Bin1^+^ OLs could be found near, but not within the 6E10^+^ amyloid deposits. As a result, the colocalisation of the Bin1 and 6E10 signals was minimal. Using six antibodies as probes of western blots (2F11, CST13679, 99D, G10, ab185950 and 1H1; for antibody target site see Fig. S1), we found a distinct expression pattern for Bin1:L and Bin1:H. Antibody clone 2F11 detected Bin1:L isoforms only, antibody clone 99D detected Bin1:H isoforms only and antibody clones CST13679, ab185950 and 1H1 detected both Bin1:H and Bin1:L isoforms (Fig. 5D). [Note: as the epitope was poorly conserved (a.a. 421-520) between human and mouse, antibody clone G10 did not detect any Bin1, as predicted]. The repeated measure analysis showed that no significant difference in Bin1:H protein levels between WT and APP/PS1, while the Bin1:L protein was significantly increased in APP/PS1, particularly the lighter 54 kDa isoform (Fig. 5E). These differences were also seen at mRNA level with a significant upregulation of Bin1:L isoforms (Fig. 5F). These results are in agreement our human findings (Summary in Table 3). In both human fAD and APP/PS1 mice, there was minimal physical association between amyloid aggregation and BIN1 expression, and no upregulation of BIN1 was observed in the areas surrounding misfolded protein. As a known inhibitor of c-Myc (Sakamuro et al., 1996), we asked if the changes in BIN1 in APP/PS1 was associated with c-Myc and found a significant increase (Fig. 5G).

**Figure 5.**
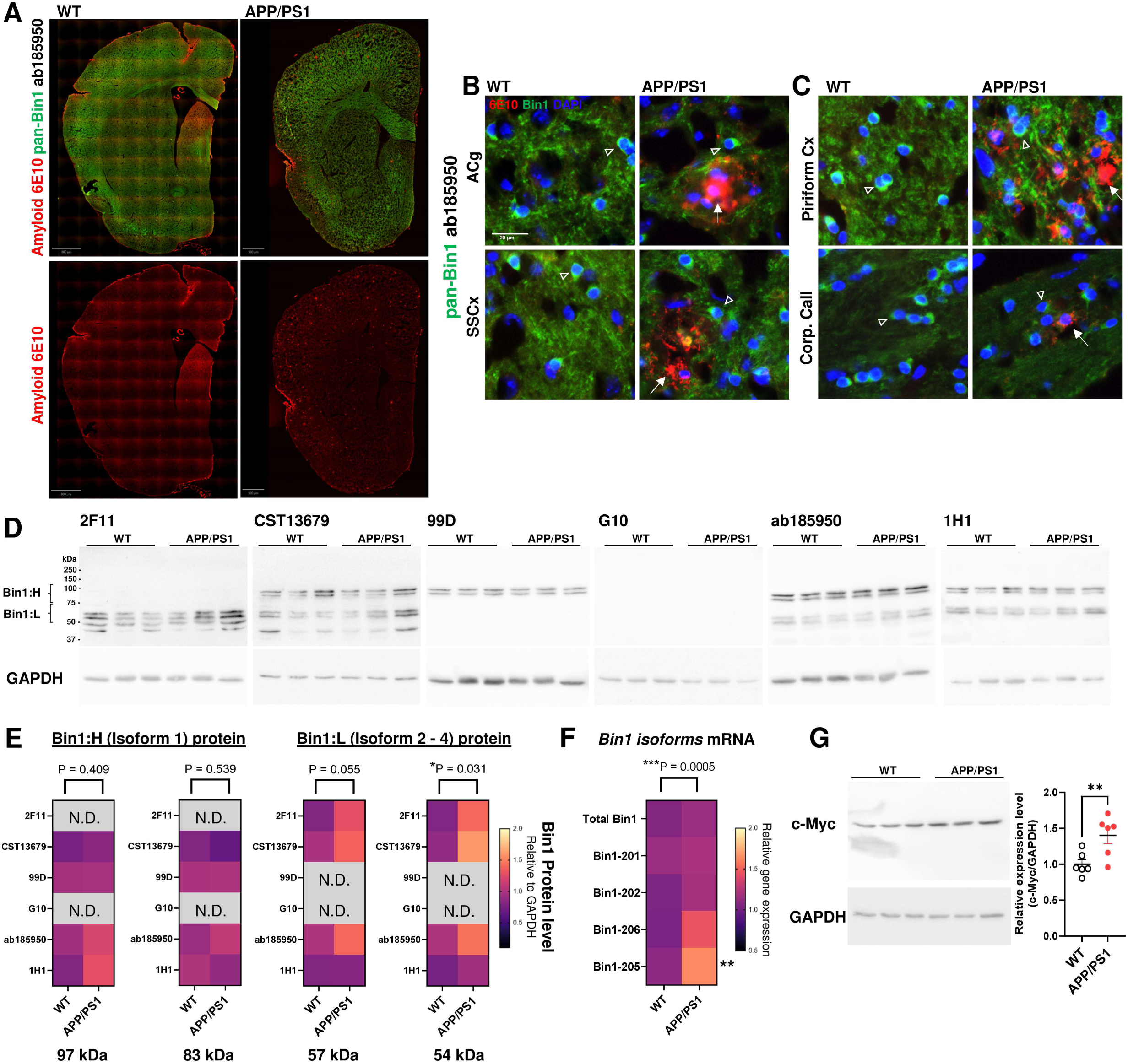
Divergent Bin1 isoforms expression in old WT and APP/PS1 mouse brains. **A** Representative whole slide immunofluorescence image of pan-Bin1 antibody (ab185950) and amyloid plaques (6E10) in 18-month-old WT and APP/PS1 brain tissues at Bregma 0.8 (Red, 6E10; Green, Bin1). **B, C** At high power, Bin1^+^ cells (open arrowheads; green; DAPI, blue) were identified across GM of anterior cingulate (ACg), somatosensory cortex (SSCx), piriform cortex (Piriform Cx) and the WM of corpus callosum (Corp. Call) with minimal difference between WT and APP/PS1. Few, if any, of these Bin1^+^ cells colocalised with 6E10^+^ signals in the amyloid plaques (white arrows) found in these regions of APP/PS1, and no morphological changes were observed in Bin1^+^ cells near amyloid plaques. **D** Representative western blots of Bin1 using antibody clones 2F11, CST13679, 99D, G10, ab185950, and 1H1 showing no difference in Bin1:H (83, 97 kDa), but an upregulation of Bin1:L (54, 57 kDa) in APP/PS1 mouse brains, with GAPDH as the loading control. **E** Heatmaps summarising the quantifications of Bin1 isoforms at 97kDa, 83kDa, 57 kDa, and 54 kDa in WT and APP/PS1 samples. The expression of OL-specific isoform (Bin1:L) was significantly increased in APP/PS1 (Two-way ANOVA, Šídák’s post-hoc, Overall ANOVA P values as denoted; n = 6). Antibody G10 did not detect any mouse Bin1 protein as predicted, due to a species difference between human and mouse (See Fig. S1) **F** Heatmaps summarising Bin1 transcript level in WT and APP/PS1 samples were Bin1:L were significantly increased in APP/PS1 (Two-way ANOVA, Šídák’s post-hoc, ** P<0.01). **G** Representative western blot of c-Myc with GAPDH as the loading control. A significant increase of c-Myc was detected in APP/PS1 (unpaired t-test ** P < 0.01; n = 6).

**Table 3.** Summary of BIN1 isoform expression pattern in human AD and APP/PS1 mouse brain.

| Anti-BIN1 |  | pan-BIN1 ab185950 |  | 99D |  | 2F11 |  |
| --- | --- | --- | --- | --- | --- | --- | --- |
| Epitope Target |  | SH3 domain |  | MBD |  | N-BAR |  |
| Human BIN1 isoform(s) Target |  | All isoforms |  | Isoform 1-3, 5-7, 9 |  | Isoform 5-7, 9,10,13 |  |
| Mouse Bin1 isoform(s) Target |  | All isoforms |  | Isoform 1, 2, X18, X20 |  | Isoform 2, X18, X20, X22, truncated |  |
| Condition / BIN1 isoforms |  | BIN1:H | BIN1:L | BIN1:H | BIN1:L | BIN1:H | BIN1:L |
| Human | Alzheimer's disease | Decrease | N.C. | Decrease | N.C. | N.D. | Increase |
| Mouse | APP/PS1 | N.C. | Increase | N.C. | N.D. | N.D. | Increase |

### Nuclear Bin1 expression in oligodendrocytes

We next validated these *in vivo* findings using in primary cultures of mouse OLs (Fig. 6A-C). Using the pan-Bin1 antibody ab185950, we found Bin1 immunostaining in both Olig2^+^ OL and MAG^+^ mOL (Fig. 6D). The majority of the immunoreactivity was localised in the cytoplasm (Fig. 6E; 26.1-26.9%; P<0.0001). Little, if any, Bin1-IR was observed in non-OL cells (Fig. 6A, E). In contrast, antibody clone 99D detected a majority of nuclear Bin1 expression among MAG^+^ mOL, NG2^+^ OPC, and non-OL cell types (NG2^-^ and MAG^-^), resembling astrocytes (Fig. 6B, F). In both NG2^+^ OPC and in MAG^+^ mOL, the proportion of nuclear Bin1 positive population was significantly higher than that in the cytoplasm (P<0.0001). Nearly every NG2^+^ OPC expressed Bin1 in the nucleus. Immunocytochemistry with antibody 2F11 that only detects Bin1:L isoforms primarily labelled OL in the cytoplasm, but not nucleus (P = 0.0187) in both NG2^+^ OPC and in MAG^+^ mOL (Fig. 6C, G). Together, the data indicate that Bin1 isoforms are found in both cytoplasm and nucleus of cells from the OL lineage and that each isoform may have its own distinct localisation pattern.

**Figure 6.**
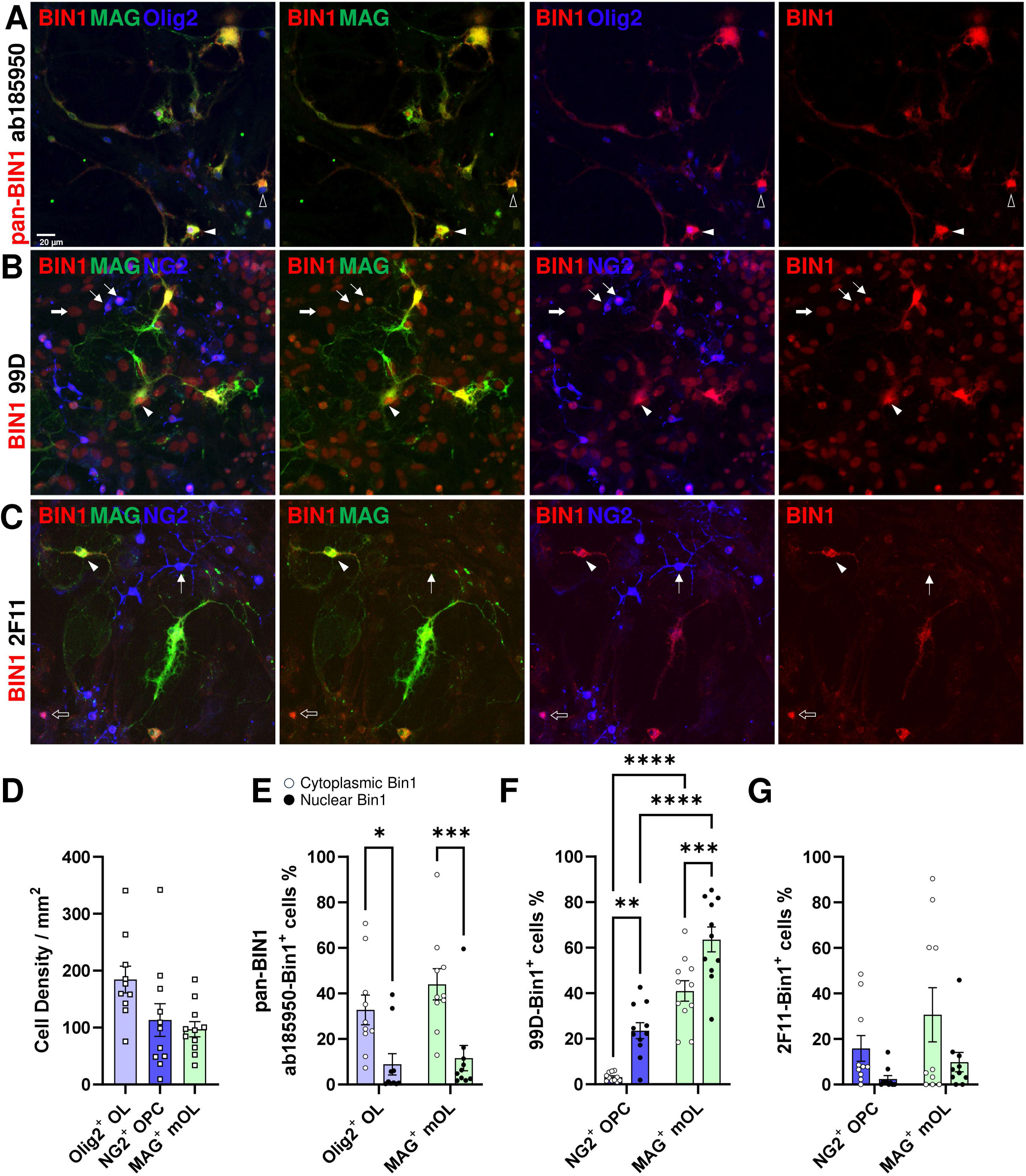
Nuclear and cytoplasmic Bin1 expression in primary murine oligodendrocyte culture. **A-C** Representative immunocytochemistry-immunofluorescence images of Bin1 expression in primary mouse OL culture detected pan-Bin1 (ab185950), 99D and 2F11 antibodies clones. **A** Bin1 (ab181950) expressions were identified in the nucleus (open arrowhead) and cytoplasm (white arrowhead) of myelinating mature OLs with colocalisation to Olig2^+^ nucleus (blue) and MAG^+^ cytoplasm (green), respectively. **B** Bin1 (99D) expressions were predominantly nuclear in NG2^+^ OPCs (blue; white arrows), MAG^+^ mature OLs (green; white arrowheads) and astrocytes with large nucleus (thick white arrow). In MAG^+^ mature OLs, Bin1 (99D) was also found in the cytoplasm or cell body. **C** Bin1 (2F11) expressions were concentrates in the cytoplasm of MAG^+^ mature OLs (green; white arrowheads). The majority of NG2^+^ OPCs (blue; open arrows) were negative for Bin1 (2F11), but few were identified (open arrow). **D** Cell density quantifications of total OL (Olig2^+^), OPC (NG2^+^) and mature OL (MAG^+^) in a typical culture. **E** Quantifications of nuclear and cytoplasmic Pan-Bin1 (ab185950) in total OL (Olig2^+^) and mature OL (MAG^+^) showing a significantly higher cytoplasmic localisation. F Quantifications of nuclear and cytoplasmic Bin1 (99D) in OPC (NG2^+^) and mature OL (MAG^+^) showing a significantly higher nuclear localisation. F Quantifications of nuclear and cytoplasmic Bin1 (2F11) in OPC (NG2^+^) and mature OL (MAG^+^) showing a higher cytoplasmic localisation. (Two-way ANOVA, Šídák’s post-hoc, *P < 0.05; **P < 0.01; ***P < 0.001; ****P < 0.0001; n = 10-12)

### Silencing Bin1 in OL perturbs pathways in both the nucleus and cytoplasm

To study the putative Bin1 functions in a purified OL model, we used siRNA to knock down Bin1 in Oli-Neu cell. This siRNA pool equally targets four regions of the *Bin1* (at positions: 251-269; 351-369; 842-860 and 1847-1865 nt). Oli-Neu is a mouse OL model cell line with endogenous c-Myc activity (Magri et al., 2014) that was generated by immortalizing mouse OPCs with the t-neu receptor tyrosine kinase overexpression (Jung et al., 1995). Immunocytochemistry and real-time PCR confirmed that siBin1 significantly decreased Bin1 protein and transcripts, compared with a scramble siRNA control (siNT, Fig. 7A) (Fig. 7B). We then performed a bulk mRNA sequencing (RNA-seq) to investigate the effect of Bin1 knockdown on the transcriptome. The quality control of the RNA-seq in principal component analysis (PCA) plot is shown in Fig. 7C (summary data in Table S5). We found that OL-specific Bin1 transcripts encoding BIN1:L protein isoforms (mouse Ensemble transcripts ID: Bin1-202, Bin1-206 and Bin1-205) were significantly reduced, while other Bin1 transcripts (Bin1-203, Bin1-204 and Bin1-207) were undetectable (Fig. 7D). As expected, the neuron specific BIN1:H transcript (Isoform 1; Ensemble transcripts ID: Bin1-201) was undetectable either by real-time PCR or RNA-seq (Fig. 7B, D). The silencing effect was similarly found across all the Bin1 exons (Fig. 7D). Among the 57,187 genes annotated, 14,061 genes were within the detection limit. Of these, 3,747 genes qualified as differential expressed (DGE, adj P < 0.05; Fig. 7E)

**Figure 7.**
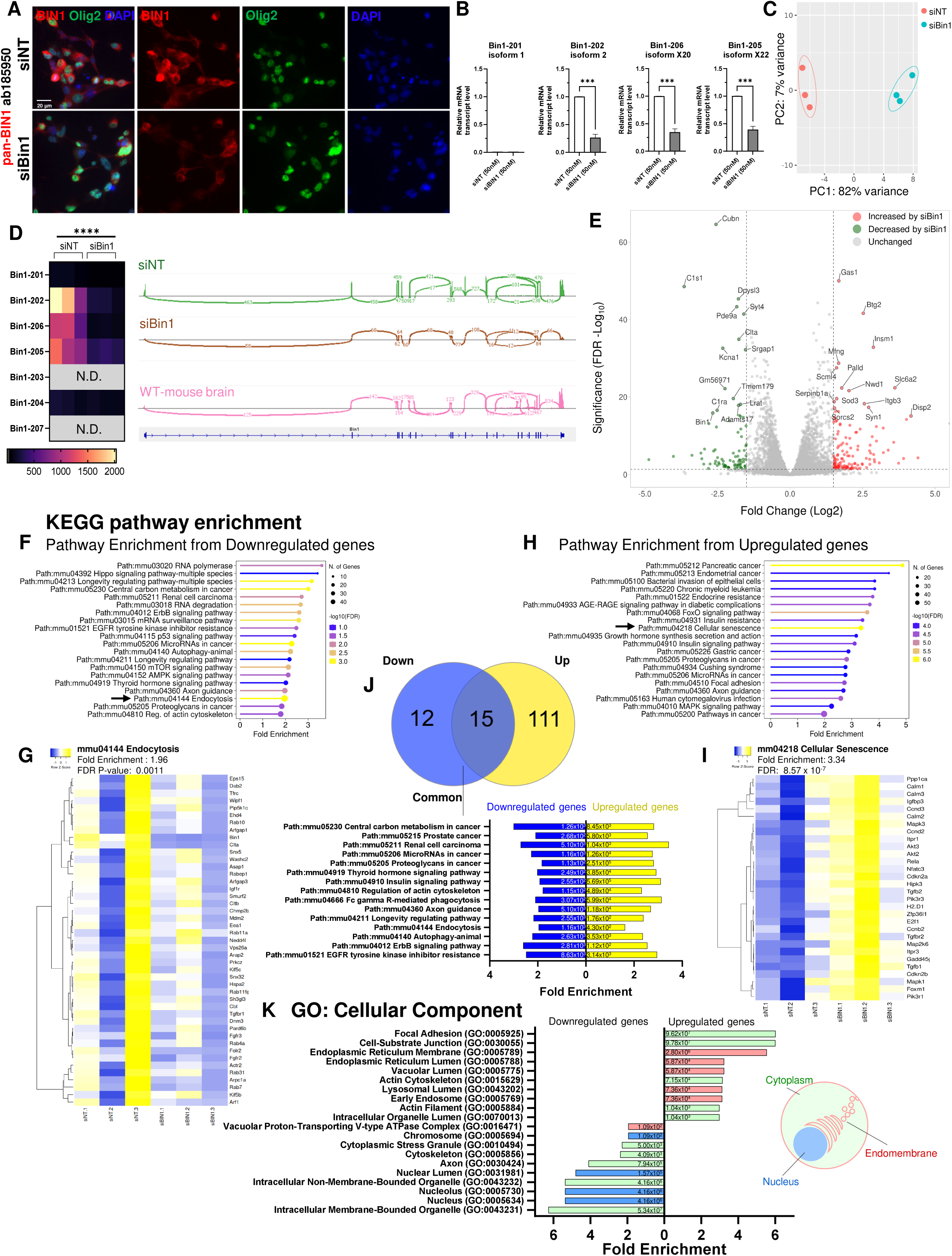
Transcriptomic shift in the OPC model upon Bin1 gene silencing. **A** Representative immunofluorescence image of pan-Bin1 (ab185950; red), Olig2 (green), and DAPI (blue) in the Oli-Neu cell culture treated with siNT and siBin1. **B** qPCR-based quantification demonstrating the effects of siBin1 on the transcription of mouse Bin1 isoform 2, Bin1 isoform 3, and Bin1 isoform 4 in Oli-Neu; Bin1 isoform 1 is not detectable (unpaired t-test vs siNT (non-target control); *** P < 0.001). **C** Principal component analysis of RNA-seq between Oli-Neu treated by siBin1 and siNT. **D** Heatmap of all Bin1 transcript variants and the sashimi plot of the RNA-seq data showing the effects of gene silencing on transcript regulation. Bin1 transcript plot from WT mouse brain provide as comparison. **E** Volcano plot of differentially expressed genes (DEGs) highlighting the top 15 significantly up-and down-regulated gene expression upon Bin1 silencing (FDR < 0.05; Fold Change >2). **F** KEGG pathway analysis of genes significantly downregulated by siBin1 in Oli-Neu **G** Endocytosis (Path: mmu04144) identified as the most significantly enriched pathway with the largest number of affected genes in F. **H** KEGG pathway analysis of genes significantly upregulated by siBin1 in Oli-Neu **I** Cellular Senescence (Path: mmu04218) identified as the most significantly enriched pathway with the largest number of affected genes in H. **J** Venn diagram of all altered pathways in F and H showing 15 common pathways identified and listed in the bar chart below. **K** Gene Ontology (cellular component) gene set enrichment confirmed that the DEGs are found both in the nucleus and the cytoplasm of Oli-Neu, with colour code shown in the legend.

We used KEGG analysis to identify the top 20 pathways enriched by the significantly down-and up-regulated genes (Fig. 7F, H). The pathway with the largest number of significantly downregulated genes were Endocytosis (Path:mmu04144;-log_10_ FDR = 2.937; nGene = 45/264). The pathway with the largest number of up-regulated genes was Cellular Senescence (Path:mmu04218;-log_10_ FDR = 6.067; nGene = 30/175) (Fig. 7F-I; Table S5). Among the 27 and 126 pathways significantly enriched by the down-and up-regulated genes, there were 15 pathways were found in common (Fig. 7J). Further analysis with gene ontology: cellular component revealed that of the top ten cell components enriched by the siBin1-downregulated genes, four were associated with nuclear functions (Fig. 7K). For the siBin1-upregulated genes, five cellular components were in the endoplasmic membrane system. Therefore, the loss of Bin1 perturbs functions in both nucleus and cytoplasm.

### Bin1 loss perturbs both Myc-and p53-related pathways

To identify the most significant molecular pathway altered by siBin1, we further our analysis using the Molecular Signatures Database (MiSigDB – Hallmark) (Fig. 8A-F). Strikingly, Myc targets were the top 2 and 3 pathways significantly enriched among the downregulated genes (Fig. 8A, B). In contrast, the p53 pathway was the most significantly enriched pathway among the upregulated genes (Fig. 8C, D). Indeed, the p53 Pathway was among the four common pathways enriched by both down-and up-regulated genes (Fig. 8E). The p53 pathway (P00059) was similarly identified as one of the 28 common mechanisms significantly perturbed by siBin1 based in an independent analysis using the PANTHER database (Fig. 8F). To identify the transcription factors likely disturbed by siBin1 in Oli-Neu cells, the gene sets were further analysed by TRRUST v2 and Enrichr-PPI based transcription factor enrichment analysis. Once again, p53 was identified by both transcription factor databases, and was in fact the only common element in the two analyses enriched by both down-and up-regulated genes (Fig. 8G). In the grid network of transcription factors, CTNNB1 (β-catenin 1) was directly associated with p53 in both up-and down-regulated genes, while Myc was directly associated with p53 in down-regulated gene (Fig. 8H). These observations suggested that the p53 signalling pathway (KEGG mmu04115), which is central to cell cycle progression, cellular senescence, DNA repair and metabolism, is a likely target of Bin1 impairment (Fig. 8I)

**Figure 8.**
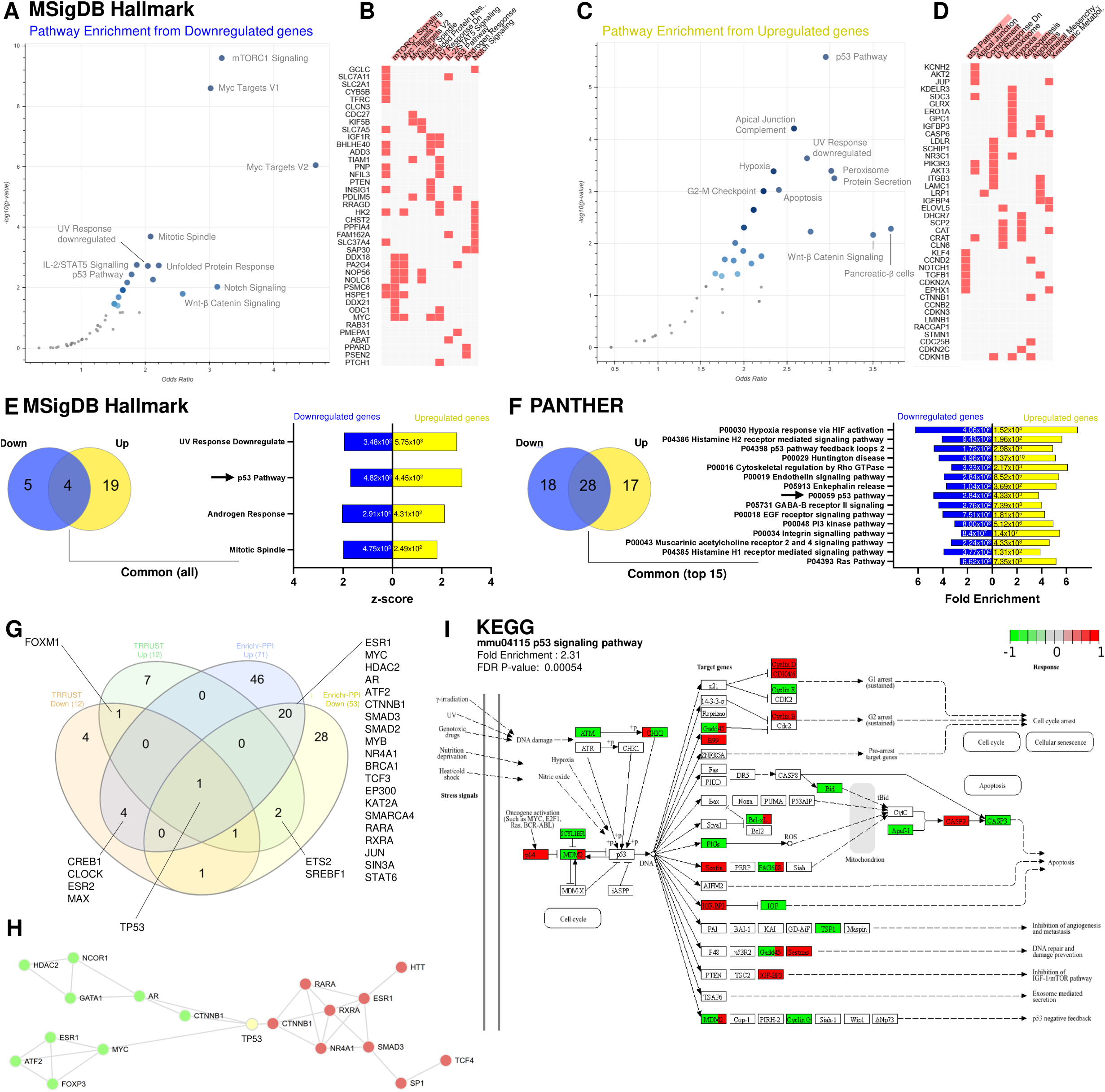
p53 signalling pathway in OPC model is perturbed after Bin1 gene silencing. **A-D** Volcano plot of gene set enrichment analysis with Molecular Signatures Database (MSigDB) hallmark database (Castanza et al., 2023) showing the significance (-log(p-value)) versus odd ratio (0, inf) based on the **A** downregulated DEGs and **C** downregulated DEGs, as reported in Fig. 7. Larger blue points represent significant terms (p-value < 0.05); smaller grey points represent non-significant terms. The intensity of blue colour correlates to the significance level. The top 3 gene sets in A were mTORC1 signalling, Myc Targets V1 and Myc Targets V2, with the top corresponding genes show on B. The top 3 gene sets in C were p53 Pathway, apical junction complement, and UV response downregulated, with the top corresponding genes show on D. **E** Venn diagram showing four common gene sets enriched in both top up-and down-regulated DEGs. **F** An independent gene set enrichment was performed with PANTHER (Protein Analysis THrough Evolutionary Relationships) (Thomas et al., 2022) as in A-D. Venn diagram showing 28 common gene sets enriched in both top up-and down-regulated DEGs with the top 15 significant enrichment listed in bar chart. **G** Venn diagram showing results of transcription factors enrichment from the up-and down-regulated DEGs in Enrichr PPI-Transcription factor (Chen et al., 2013) and TRRUST v2 (Han et al., 2018) databases. Transcription factors enriched by down-regulated DEGs in TRRUST v2 (orange), Up-regulated DEGs in TRRUST v2 (green), Down-regulated DEGs in Enrichr PPI (blue), Up-regulated DEGs in Enrichr PPI (yellow) were shown. The common transcription factors perturbed by each enrichment are listed.TP53 (p53) is the only transcription factor commonly perturbed by all four enrichment. **H** The TP53 centred grid map based on the Enrichr PPI-Transcription factor enrichment showing the interacting protein-protein interaction network affected by TP53 mediated by the downregulated (green) and upregulated (red) DEGs mediated by siBin1. **I** KEGG diagram of mouse p53 signalling pathway (mmu04115) mapped with all the significant upregulated (red) and downregulated (green) DEGs mediated by siBin1 in the OPC model.

### The loss of nuclear BIN1 disinhibits cell cycle progression

BIN1 is a known c-Myc inhibitor and tumour suppressor that reduces cell cycle progression in OL (Magri et al., 2014). In STRING-based protein-protein interaction analysis, c-Myc connects BIN1 with p53 indirectly in both human and mouse (Fig. 9A). We therefore hypothesised that the perturbation of p53 pathways and the altered cell cycle events subsequent to Bin1 loss were mediated by c-Myc (Fig. 9B). To identify the Bin1 isoform responsible for this effect, we performed immunocytochemistry on Oli-Neu cells treated with siBin1 using pan-Bin1 (ab185950), 99D and 2F11 antibodies (Fig. 9C). We found that the pan-Bin1-IR (ab185950), was significantly reduced in both cytoplasmic and nuclear regions (Fig. 9D, E). In contrast, antibody clone 99D detected a significant reduction of cytoplasmic Bin1, but not in the nucleus (Fig. 9F, G). In addition, the Bin1:L specific antibody, 2F11, was not significantly altered in either the cytoplasm or nucleus after the genetic knockdown (Fig. 9H, I). Intriguingly, siBin1 had no discernible effects on overall cell density, Olig2^+^ nuclei density nor c-Myc expression level (Fig. 9J). Despite this, the proportion of cells that were engaged in cell cycle activity (Cyclin D1^+^ Olig2^+^ double positive population) were significantly increased. As Cyclin D is one of the targets downstream to c-Myc-mediated cell cycle progression (Dong et al., 2014), this finding suggested that Bin1 loss may disinhibit c-Myc activity in OL.

**Figure 9.**
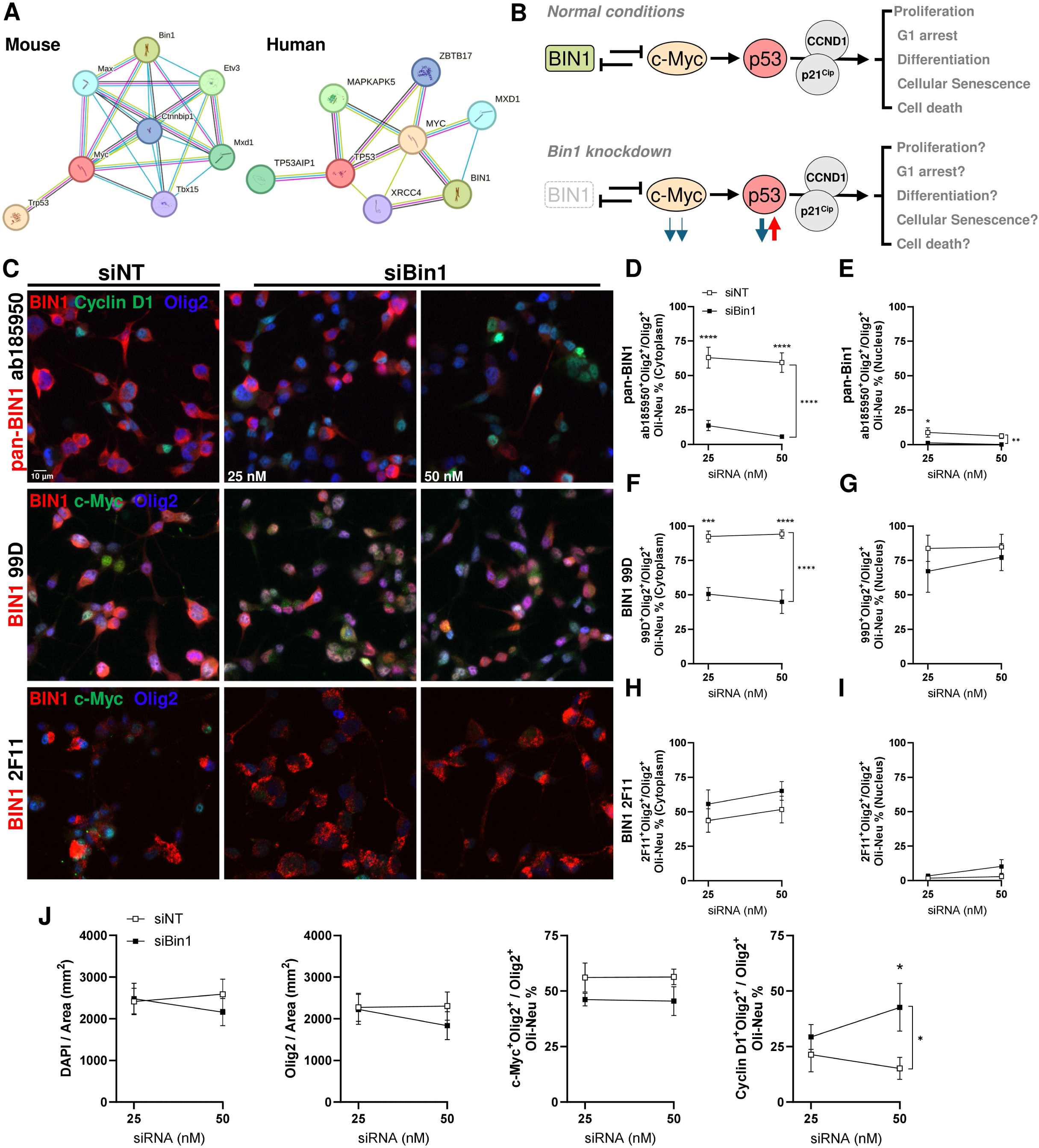
**Deficiency in BIN1**-**c-Myc interactions disrupt cell cycle regulation A** Protein-protein interaction network results from a prompt of BIN1, c-Myc and p53 in human and mouse using STING functional protein association networks analysis. BIN1 has indirect, but no direct, association with p53 via Myc in both human and mouse. The evidence are indicated by the coloured lines: purple, experimental evidence; green, neighbourhood evidence; blue, cooccurrence evidence; yellow, textmining evidence; light blue, database evidence; black, co-expression evidence (https://string-db.org/)**. B** Schematic diagram of RNA-seq data driven hypothesis of BIN1-c-Myc interactions-mediated p53 regulated cell cycle control in OPCs in with or without BIN1. **C** Representative immunocytochemistry-immunofluorescence images of Oli-Neu OPC model treated siBin1 (25 or 50 nM) or control (siNT) using BIN1 antibody clones ab181950 (top), 99D (middle), or 2F11 (below) with anti-Cyclin D1 (top) or anti c-Myc (middle and below). Cytoplasmic Bin1 were detected by all Bin1 antibodies. Colocalisation of Bin1 and c-Myc in the Olig2^+^ nuclei were observed in 99D. Red, Bin1; green, Cyclin D1 or c-Myc; blue, Olig2. **D - I** Quantifications of the cytoplasmic (**D** Ab185950, **F** 99D, and **H** 2F11) and nuclear Bin1 (**E** ab185950, **G** 99D, and **I** 2F11) showing a significant reduction of cytoplasmic (ab185950, 99D) and nuclear (ab185950) Bin1 signal in the Olig2^+^ Oli-Neu cells upon siBin1 treatment (25 or 50 nM, vs siNT) (Two-way ANOVA, Šídák’s post-hoc, *P < 0.05; **P < 0.01; ***P < 0.001; ****P < 0.0001; n = 4; ANOVA P value denoted between lines). **J** Quantifications of cell density (DAPI), Olig2^+^ nuclei density, c-Myc^+^ nuclei density and Cyclin D1^+^ nuclei density upon siBin1 treatment (25 or 50 nM, vs siNT) (Two-way ANOVA, Šídák’s post-hoc, *P < 0.05; **P < 0.01; ***P < 0.001; ****P < 0.0001; n = 4; ANOVA P value denoted between lines). A significant upregulation of Cyclin D1 was found in siBin1.

### Bin1-c-Myc interactions in OL lineage

To further delineate the Bin1-c-Myc relationship, we assayed changes in Bin1 isoforms in Oli-Neu model following the application of the selective c-Myc inhibitor, 10058-F4 (Fig. 10A, B). Inhibition of endogenous c-Myc activity by 10058-F4 significantly reduced pan-Bin1 (ab185950) antibody staining in the cytoplasm but did not affect nuclear Bin1 levels (Fig. 10B). However, 10058-F4 significantly increased the nuclear Bin1 expression detected by 99D in a concentration dependent fashion without an effect on cytoplasmic Bin1. By contrast, after 10058-F4 mediated c-Myc inhibition, Bin1:L immunoreactivity (2F11) was increased in both the nucleus and cytoplasm and Olig2^+^ cell density was reduced by 30%. Myc inhibition also significantly increased the proportion of c-Myc^+^ and Cyclin D1^+^ cells by 7.6-fold (Fig. 10C). Immunoblotting confirmed that c-Myc inhibition triggered the upregulation of Bin1:L isoforms at the protein (Fig. 10D, F; ab185950 and 2F11), and transcript level (Fig. 10F). No change in c-Myc protein or transcript level was detected (Fig. 10D, F, G).

**Figure 10.**
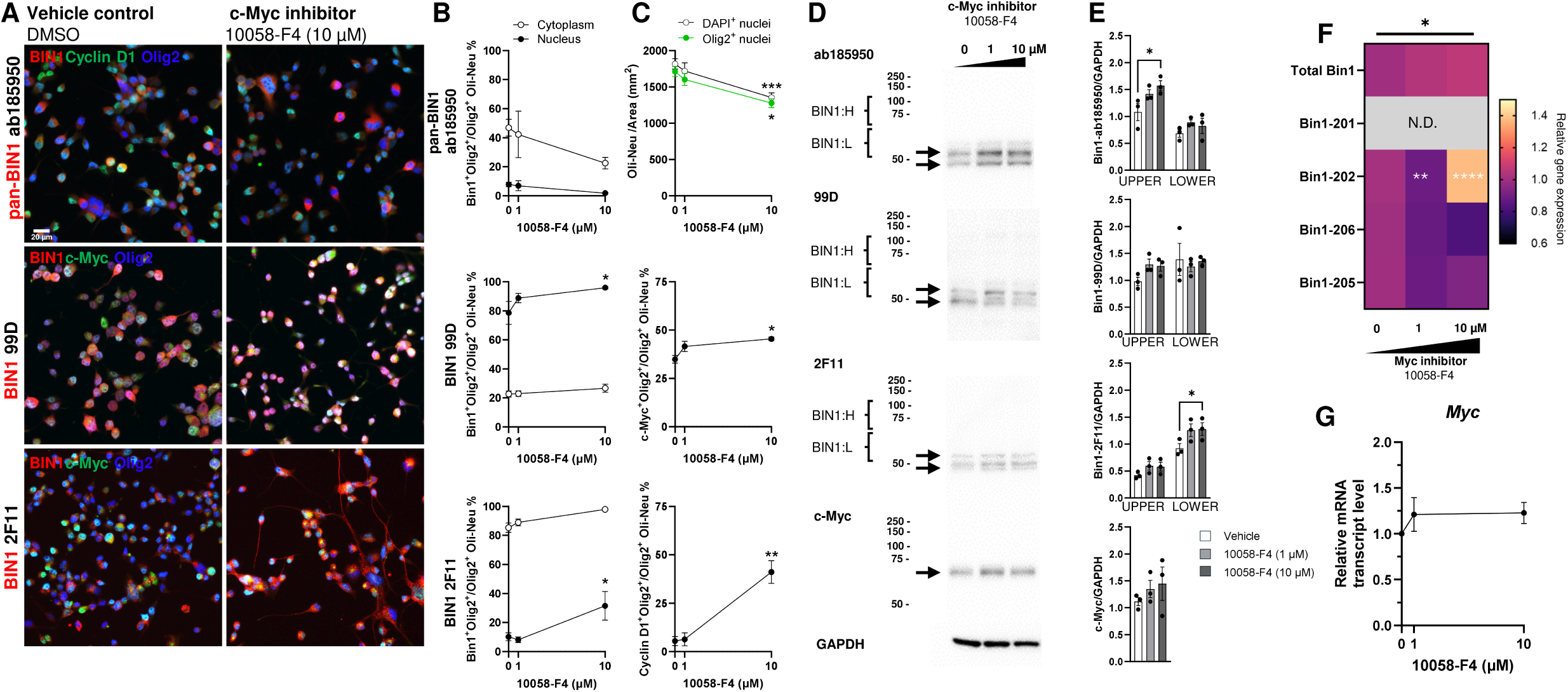
c-Myc inhibition increases Bin1:L expression and their nuclear translocation. **A** Representative immunocytochemistry-immunofluorescence images of Oli-Neu OPC model treated by c-Myc inhibitor (10058-F4, 10 µM; 24 h) or control (DMSO vehicle) using BIN1 antibody clones ab181950 (top), 99D (middle), or 2F11 (below) with anti-Cyclin D1 (top) or anti c-Myc (middle and below). Red, Bin1; green, Cyclin D1 or c-Myc; blue, Olig2. Cytoplasmic Bin1 were detected by all Bin1 antibodies. Colocalisation of Bin1 and c-Myc in the Olig2^+^ nuclei were observed, and a clear translocation of 2F11 Bin1 into nucleus was observed after c-Myc inhibition (below). **B** Quantifications of the Bin1 using antibody clones ab185950 (top), 99D (middle), and 2F11 (below) in the cytoplasm (open circle) and nuclei (black circle) as in A. A significant increase of 99D and 2F11 in the nucleus, but not cytoplasm was detected (Two-way ANOVA, Šídák’s post-hoc, *P < 0.05 vs DMSO; n = 3). **C J** Quantifications of cell density (DAPI) and Olig2^+^ nuclei density (green circle)(top), c-Myc^+^ nuclei density (middle) and Cyclin D1^+^ nuclei density (below) treated by c-Myc inhibitor (10058-F4, 1 or 10 µM) or control (DMSO vehicle). A significant reduction of cell density with a significant increase of nuclear c-Myc and Cyclin D1 were found. (Two-way ANOVA, Šídák’s post-hoc, *P < 0.05; **P < 0.01; ***P < 0.001; vs DMSO; n = 3). **D** Representative western blots of Bin1 in Oli-Neu OPC model treated by c-Myc inhibitor (10058-F4, 10 µM) or control (DMSO vehicle) using antibody clones ab181950, 99D or 2F11, and of c-Myc with GAPDH as the loading control. **E** Quantification of Bin1 or c-Myc expression in D where 10058-F4 mediated c-Myc inhibition had an overall significant effect on Bin1 expression detected by of pan-Bin1 antibody (ab185950) (P = 0.0246) and 2F11 (P = 0.0221), with pairwise difference in upper (57 kDa) and lower (54 kDa) bands indicated on graphs (Two-way ANOVA, Šídák’s post-hoc, *P < 0.05; **P < 0.01; ***P < 0.001; vs DMSO; n = 3). No significant differences were found in 99D or c-Myc (One-way ANOVA, Tukey post-hoc; n = 3). **F** Heatmaps summarising *Bin1* transcript level in Oli-Neu OPC model treated by c-Myc inhibitor (10058-F4, 10 µM; 24 h) or control (DMSO vehicle). 10058-F4 had an overall significant effect on *Bin1* transcription (P = 0.042) and Bin1-202 transcript (isoform 2) was significantly changed in a concentration-dependent fashion. (Two-way ANOVA, Šídák’s post-hoc, *P < 0.05; **P < 0.001; ****P < 0.0001; n = 3). **G**. No changes was found in *Myc* transcript level upon 10058-F4 inhibition. N.D. Not detectable

### Bin1-c-Myc-Cyclin D expression during oligodendrocyte differentiation

Next, to determine if Bin1-mediated c-Myc inhibition was required for OL differentiation, we examined the myelinating phenotype of Oli-Neu cells after Bin1 knockdown. Although siBin1 exerted no effects on the levels of MAG (Fig 11A), it significantly reduced the transcription of the master myelin transcription factor, *Myrf* (Fig.11B). Indeed, gene ontology analysis of the RNA-seq dataset confirmed that myelination (GO:0042552) was one of the most significantly disrupted biological process affected by Bin1 knockdown in Oli-Neu cells (Fig. 11C). To address the effects of differentiation, we compared the levels of Bin1 isoforms in Oli-Neu cells cultured for 48 hours in either growth media (GM) or differentiation media (DM) (Fig. 11D). The differentiation media had no effects on pan-Bin1 expression (ab185950) or immunostaining of Bin1 isoform 1-2 specific antibody (99D) (Fig. 11E, F). However, differentiation significantly reduced both the cytoplasmic and nuclear Bin1:L levels detected by 2F11 (P = 0.0057). Phenotypically, differentiation did not alter the cell density or the proportion of c-Myc^+^ cells, but it significantly reduced cycling (Cyclin D1^+^) cells by 25.9% (Fig. 11G). Differentiation did not alter the levels of Bin1 protein isoforms (Fig. 11H), but significant reductions of Bin1 and Myc mRNA were found (Fig. 11I, J). The changes of BIN1 protein isoforms expression in OLs culture experiments are summarised in Table 4.

**Figure 11.**
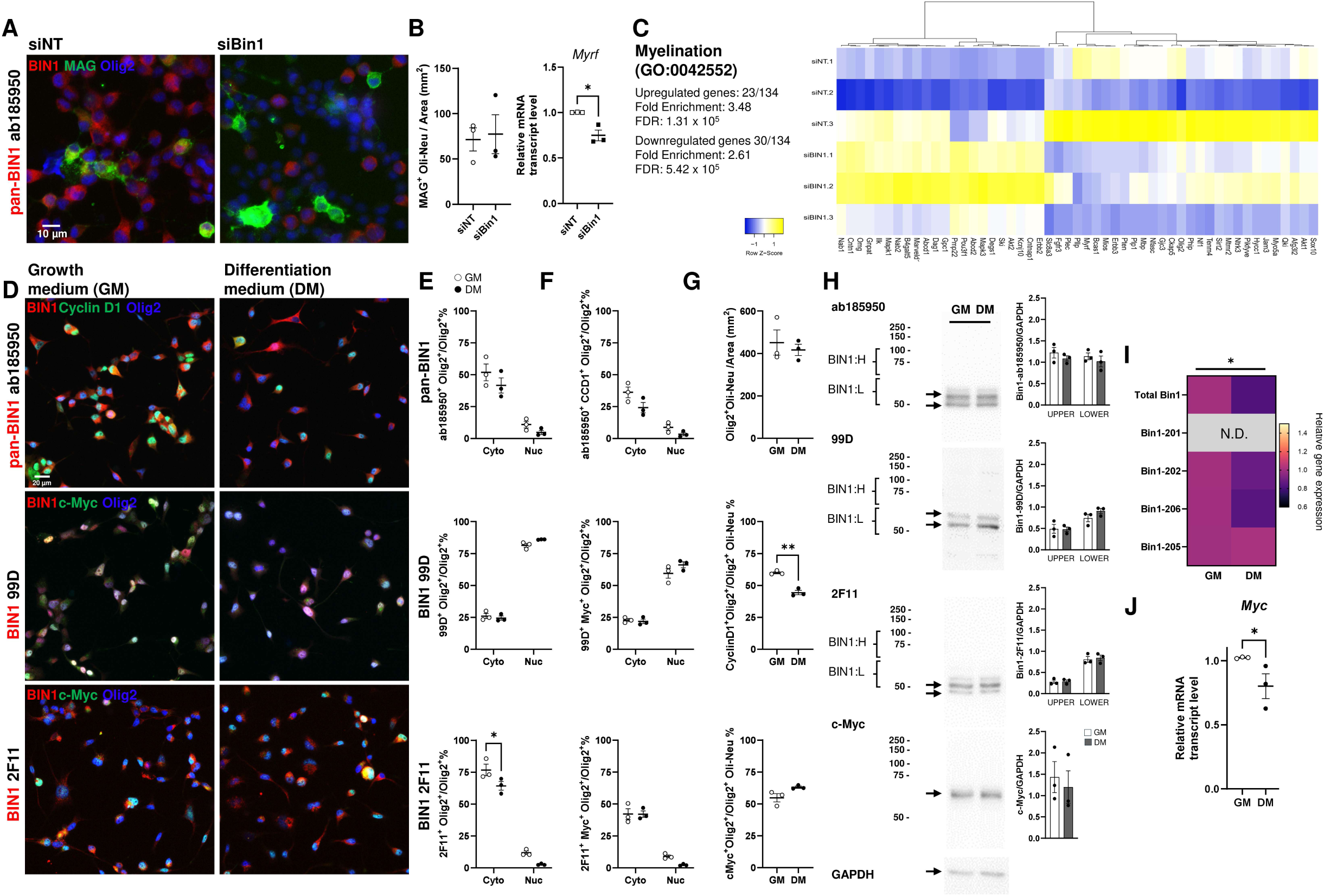
Differentiation of oligodendrocyte precursor cells does not affect Bin1 expression profile and localisation. **A** Representative immunocytochemistry-immunofluorescence images of Oli-Neu OPC model after siBin1 silencing with pan-Bin1 (ab185950; red), MAG (green), and Olig2 (blue). **B** Quantifications of immunocytochemistry showing no significant differences in MAG^+^ mOL formation between siNT and siBin1, but a significant reduction of *Myrf* mRNA transcript was found (unpaired t-test, *P < 0.05; n = 3). **C** Myelination process (GO:0042552) was significantly enriched in the RNA-seq data (see Fig. 7) in the Oli-Neu OPC model after Bin1 silencing. **D** Representative immunocytochemistry-immunofluorescence images of Oli-Neu OPC model in proliferative state with growth medium (GM) or after differentiation media (DM) incubation for 48 hours. BIN1 antibody clones ab181950 (top), 99D (middle), or 2F11 (below) with anti-Cyclin D1 (top) or anti c-Myc (middle and below) are shown. Red, Bin1; green, Cyclin D1 or c-Myc; blue, Olig2. Nuclear Cyclin D1 was remarkably reduced. **E** Quantifications showed a minimal effect of differentiation (DM) on Bin1 expression or their nucleocytoplasmic translocation of ab185950 or 99D but had an overall significant effect on 2F11 (P = 0.0057) (Two-way ANOVA, Šídák’s post-hoc, *P < 0.05 GM vs DM; n = 3). **F** Quantifications showed a minimal effect of differentiation (DM) on Cyclin D1^+^ Bin1^+^ or c-Myc^+^ Bin1^+^ double positive cells among all antibody clones. **G** Quantifications of Olig2^+^ cell density (top), Cyclin D1^+^ (middle) and c-Myc^+^ nuclei (below) density showing a significant reduction of Cyclin D1^+^ cells upon differentiation (unpaired t-test, **P < 0.01; n = 3). **H** Representative western blots of Bin1 in Oli-Neu OPC model upon differentiation (DM) using antibody clones ab181950, 99D or 2F11, and of c-Myc with GAPDH as the loading control. Quantifications of these blots of Bin1 are shown on right but no significant differences were found (Bin1: Two-way ANOVA, Šídák’s post-hoc, *P < 0.05; **P < 0.01; ***P < 0.001; vs DMSO; n = 3; c-Myc: One-way ANOVA, Tukey post-hoc; n = 3). **I** Heatmaps summarising *Bin1* transcript level in Oli-Neu OPC model upon differentiation (DM) showing an overall significant effect on *Bin1* transcription (P = 0.0329) (Two-way ANOVA, Šídák’s post-hoc; n = 3). **J**. RT-PCR results showing a significant reduction of *Myc* transcript upon differentiation (unpaired t-test, *P < 0.05; n = 3). N.D. Not detectable

**Table 4.**
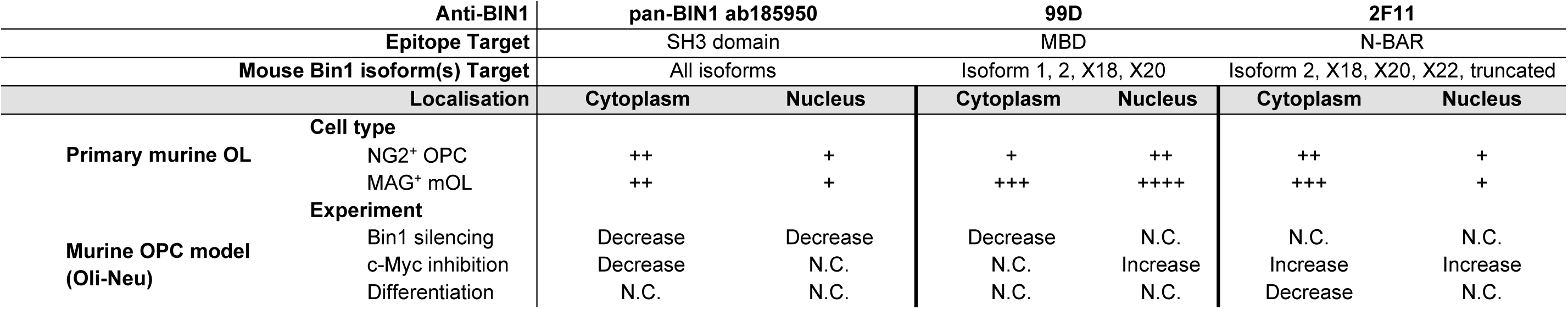
Summary of BIN1 isoform expression in OL culture.

| Anti-BIN1 |  | pan-BIN1 ab185950 |  | 99D |  | 2F11 |  |
| --- | --- | --- | --- | --- | --- | --- | --- |
| Epitope Target |  | SH3 domain |  | MBD |  | N-BAR |  |
| Mouse Bin1 isoform(s) Target |  | All isoforms |  | Isoform 1, 2, X18, X20 |  | Isoform 2, X18, X20, X22, truncated |  |
| Localisation |  | Cytoplasm | Nucleus | Cytoplasm | Nucleus | Cytoplasm | Nucleus |
| Primary murine OL | Cell type |  |  |  |  |  |  |
|  | NG2<sup>+</sup> OPC | ++ | + | + | ++ | ++ | + |
|  | MAG<sup>+</sup> mOL | ++ | + | +++ | ++++ | +++ | + |
| Murine OPC model<br>(Oli-Neu) | Experiment |  |  |  |  |  |  |
|  | Bin1 silencing | Decrease | Decrease | Decrease | N.C. | N.C. | N.C. |
|  | c-Myc inhibition | Decrease | N.C. | N.C. | Increase | Increase | Increase |
|  | Differentiation | N.C. | N.C. | N.C. | N.C. | Decrease | N.C. |

### BIN1:H and BIN1:L isoforms bind to c-Myc in silico

The regulation BIN1:L isoforms expression in human brain tissues, mouse brain tissues and OL cell culture were distinct from BIN1:H in disease and experimental conditions. To predict the extent to which various BIN1:H and BIN1:L isoforms to interact with c-Myc (Fig. 12A), we used ClusPro and AlphaFold database and ClusPro to screen all the available models of human and mouse BIN1 isoforms for the probability of their docking with the corresponding models of c-Myc (Table S6). Among the six successful docking experiments between human BIN1 and c-Myc, there were no significant differences in the putative affinity (lowest energy of hydrophobic interactions and members involved) between BIN1:L (isoforms1 - 3) and BIN1:H (isoforms 9,10, 13) (Fig. 12B, C). An earlier study reported a potential binding site between the SH3 domain of human BIN1 peptide and the MB1 (Myc Box 1) sequence of a c-Myc fragment (Pineda-Lucena et al., 2005) (Fig 12D). We have now extended this work and validated the interaction between BIN1 and c-Myc at the SH3 and MB1 region using the whole protein model of BIN1:H (isoform 1) (Fig. 12E). More importantly, a highly similar interaction was identified between BIN1:L (isoform 9) where the SH3 domain was strongly associated with the MB2 (Myc Box 2) region of human c-Myc (Fig. 12F), a key determinant of c-Myc transcriptional activity (Andresen et al., 2012).

**Figure 12.**
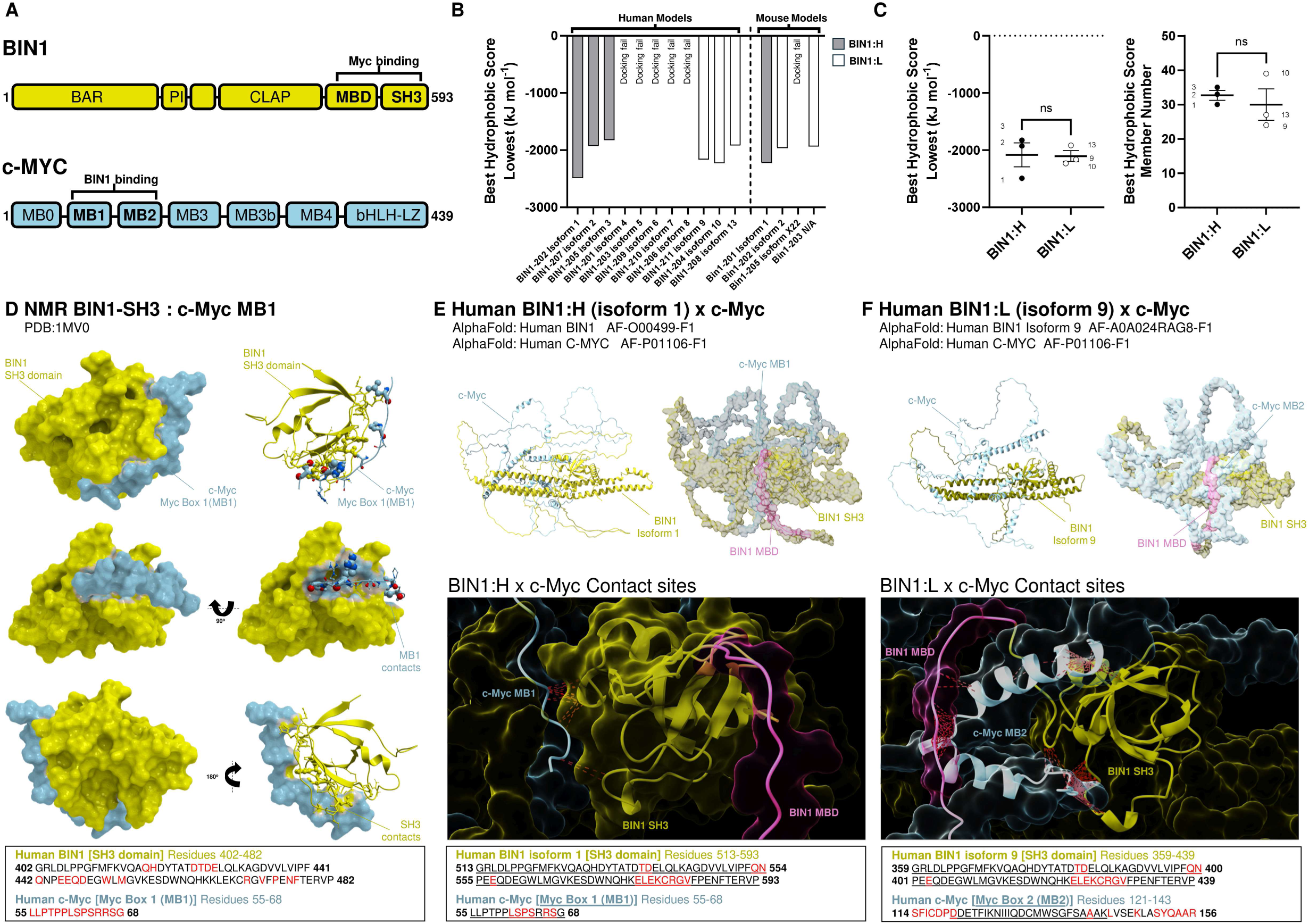
Human BIN1:L and BIN1:H similarly binds to c-Myc at MB1 and MB2 *in silico*. **A** Schematic diagram of the BIN1 and c-MYC structure of human origin. Function domains are annotated. The reported Myc and BIN1 binding sites are shown. MB: Myc Box; bHLH-LZ: Basic Helix-Loop-Helix Leucine Zipper domain **B** Bar chart showing the in silico docking results (ClusPro 2.0 protein-protein docking server (Kozakov et al., 2017)) between all available BIN1 structures in AlphaFold database and c-Myc of human and mouse origin respectively. The lower the energy level (Best Hydrophobic Score Lowest kJ mol^-1^) represents the stronger binding affinity. Details of the models in Table S6 **C** Dot plot showing comparison between BIN1:H (isoform 1-3) and BIN1:L (isoform 9, 10, 13) in the binding affinity and the numbers of residues in the putative binding site (unpaired t-test; n = 3; Isoform numbers are indicated adjacent to each data point)). **D** *Left* The reported experimental binding model (PDB: 1MV0; (Pineda-Lucena et al., 2005)) between human BIN1-SH3 (yellow) and c-Myc-MB1 (blue), with 90° (Y axis) and then 180° (Z axis) rotation. *Right* Corresponding model with either BIN1-SH3 and/or c-Myc-MB1 presented as ribbons and residues. All contact area at MB1 and SH3 are highlighted on the skin with reciprocal colours. The sequences in the binding structures are shown in the box below, with residues in close contact in red font. **E** Human BIN1:H (yellow) and c-Myc (blue) binding representation by BIN1 isoform 1 (AlphaFold model as denoted). *Top* The known interacting sites including BIN1 SH3, BIN1 MBD (pink) and c-Myc MB1. *Middle* A close up presentation of the binding site between BIN1 SH3 and c-Myc MB1 with residue contacts connected by dotted red lines. BIN1 MBD is not in contact but at clos proximity with BIN1 SH3. *Below* The sequences in the binding model are shown in the box, with residues in close contact in red font. **F** Human BIN1:L (yellow) and c-Myc (blue) binding representation by BIN1 isoform 9 (AlphaFold model as denoted). *Top* The known interacting sites including BIN1 SH3, BIN1 MBD (pink) and c-Myc MB2. *Middle* A close up presentation of the binding site between BIN1 SH3 and c-Myc MB2, as well as between BIN1 MBD and c-Myc MB2, with residue contacts connected by dotted red lines. *Below* The sequences in the binding model are shown in the box, with residues in close contact in red font.

## DISCUSSION

The *BIN1* gene encodes a family of nucleocytoplasmic proteins whose size and structure vary based on alternative splicing of its 20 exons in human and mouse (Dourlen et al., 2025; Taga et al., 2020; Tan et al., 2013), but the subcellular localisation and physiological functions of its many isoform in OLs and other brain cells, remain largely unknown (De Rossi et al., 2016; Ponnusamy et al., 2023). The present study offers new details concerning the differential expression of BIN1 isoforms in human and mouse brain, their nucleocytoplasmic distribution in OL, and provide insight into their possible nuclear functions.

In agreement with earlier studies (Adams et al., 2016; De Rossi et al., 2016), we observe that most small non-neuronal cells in the cortex are strongly positive for BIN1. In the GM, many of these small cells express nuclear Olig2 and are identified as perineuronal satellites, and thus mostly likely are members of the OL lineage. Significantly, BIN1 immunoreactivity was found in the nucleus as well as in the cytoplasm of such cells. In WM strings of cells with elongated nuclear shapes were found with similar distributions of BIN1 in the nucleus and cytoplasm. The nuclear localisation of BIN1in human brain has been recognised for more than 20 years, (DuHadaway et al., 2003), but it has received little attention in subsequent investigations on the role of BIN1 in AD pathogenesis (De Rossi et al., 2019; De Rossi et al., 2016; Ponnusamy et al., 2023; Taga et al., 2020; Zhang et al., 2024). Given its known connection with c-Myc, the overlooking of BIN1 nuclear function is surprising. The c-Myc oncogene is a well-known driver of cell cycle activity. BIN1 was subsequently discovered to be a c-Myc interacting protein with properties of a tumour suppressor that inhibited cell cycle progression. (Cassimere et al., 2009; Elliott et al., 2000; Elliott et al., 1999; Pyndiah et al., 2011; Sakamuro et al., 1996; Tan et al., 2013). As aberrant cell cycle activity is a prominent feature of both neurons (Herrup and Yang, 2007), and myelinating OLs at risk for degeneration in AD (Tse et al., 2018), elucidating the BIN1–Myc relationship in the nucleus is of considerable importance.

*MYC* is tightly regulated in glial cells that maintain cell cycle activity (Faria et al., 2008). As OPCs are self-renewing precursors, it was not suppressing to observe the positive association between BIN1 and c-Myc expression level in the OPC model. The OPC population persists in the adult brain through self-renewal and maintains the myelinating capacity in postnatal development. In the switch between cell cycle progression and differentiation in OPCs, c-Myc was identified as the key regulator (Magri et al., 2014). c-Myc overexpression has also been shown to be the determining factor in the rejuvenation of an ageing or dormant OPC into an active OPC (Neumann et al., 2021). As myelinating OLs are postmitotic, finding the nuclear Bin1 expression in these cells was unexpected. In sporadic AD, DNA damage triggers aberrant cell cycle protein expression (Tse et al., 2018; Tse and Herrup, 2017) and abnormal cell cycle re-entry is known to cause OL cell death (Grinspan et al., 1996; Tse et al., 2018). It is plausible therefore that nuclear Bin1 expression regulate cell cycle in both OPC and postmitotic OL in health and disease by controlling c-Myc activity.

In this context, it is important to better define the relationship between BIN1 function and c-Myc activity in OLs. In Oli-Neu cells, which only expresses BIN1:L isoforms, silencing Bin1 results in the perturbation of p53 and c-Myc signalling and the disruption of cell cycle regulation. BIN1 silencing strongly upregulated Cyclin D1 without any significant cell death or proliferation. This growth arrest in the OPC model coincided with a significantly enriched pathways downstream to p53 signalling including cellular senescence. Under physiological conditions, Cyclin D1 expression is required for OPC proliferation (Bansal et al., 2005). However, Cyclin D1 is stimulated by p53 activation, and it mediates cellular senescence through p21^CIP1^ (Chen et al., 1995; Lossaint et al., 2022). These data suggested that the loss of BIN1:L to regulate c-Myc activity leads to the cellular senescence program through p53-p21^CIP1^ in OPC as reported earlier (Rivellini et al., 2022).

BIN1 variants confer significantly increased risk of AD. These studies, however, did not parse the effects of the genetic variants on the different BIN1 isoforms. Here, using specific primers against each BIN1 transcript and six antibodies against specific BIN1 structural domains among these isoforms, we have confirmed the divergent regulation of BIN isoforms in human AD brain. While GWAS-identified variants of *BIN1* in sporadic AD variants (e.g. rs754834233, [P318L]; rs138047593, [K358R]) have been shown to accelerate amyloidosis (Perdigao et al., 2021), or impair synaptic vesicle exo-endocytosis (Barata et al., 2025), the genetic constructs used in these studies were based on the neuron-specific BIN1:H isoform (isoform 1, NP_033798.1). Our finding of a significant increase in the OL-specific BIN1:L protein isoforms in both human AD and its AD models validates and extends earlier findings (De Rossi et al., 2016), that is, the increase in OL-enriched BIN1:L and along with a decrease in neuron-enriched BIN1:H isoforms in AD.

Previous studies reported the possible association between BIN1 expression and misfolded proteins accumulation in AD brain, such as tau (Chapuis et al., 2013; Ponnusamy et al., 2023; Taga et al., 2020; Zhang et al., 2024) or amyloid (De Rossi et al., 2019; Miyagawa et al., 2016; Perdigao et al., 2021). We find no evidence for the presence of Bin1 within the senile plaques found in familial AD brains, nor its colocalisation with amyloid immunoreactivity in the AD mouse model. The significant reduction of BIN1:H that contains the CLAP domain may be related to the response for misfolded protein clearance through endocytosis (Taga et al., 2020; Zhang et al., 2024), but the BIN:L isoforms significantly upregulated in AD contain minimal CLAP domain. By using specific primers and antibody clone 2F11 that targets the junction between *BIN1* exon 6 and 8 (BIN1:L isoforms only; i.e. BIN1 isoforms 5-7, 9, 10, 13 in man, isoforms 2, X20, X22 in mouse) (DuHadaway et al., 2003), and antibody clone 99D that detects any isoforms with exon 17 in the MBD domain (Both BIN1:L and BIN1:H isoforms; i.e. BIN1 isoforms 1-3, 5-7, 9 in man, isoforms 1-2 in mouse), our cell culture experiments confirmed that OL lineage only express BIN1:L isoforms. It remains unknown if the GWAS-identified *BIN1* variants impair BIN1:L isoforms in OL, as in neurons (Barata et al., 2025; Perdigao et al., 2021). Further, most of the GWAS identified variants (e.g. rs59335482, rs744373, rs6733839, rs4663105) are upstream of *BIN1* gene within the enhancer region and thus would have no direct impact on the amino acid sequences encoded by the 20 *BIN1* exons (Bellenguez et al., 2022; Chapuis et al., 2013; Franzmeier et al., 2019; Harold et al., 2009; Qiu et al., 2022; Seshadri et al., 2010).

Given our new findings, investigations into the impact of these variants on the regulation of BIN1 splicing in neurons and OLs in AD pathogenesis are clearly warranted.

Using constructs from targeted mutagenesis (Elliott et al., 1999), or small construct of BIN1 peptides (Andresen et al., 2012; Pineda-Lucena et al., 2005), early reports identified the MBD and SH3 domains of BIN1 as the specific sites of c-Myc interactions. In a docking experiment with a partial 3D model, the BIN1 SH3 domain was shown to bind Myc Box 1 at residues 55-68 of c-Myc (Pineda-Lucena et al., 2005). In agreement with a recent proteomics-based interactome analysis of BIN1 SH3 domain (Zambo et al., 2024). These authors proposed that SH3 binds to the proline-rich PxxP motif (Pro-x-x-Pro) that can be found in a large number of key proteins involved in nuclear processes and mitosis, including c-Myc. As both MB1 and MB2 are key determinant of the protein-protein interaction and transcriptional regulatory machinery in c-Myc (Llombart and Mansour, 2022), our simulation experiment suggests that BIN1:L binds to c-Myc in a fashion similar to that identified for BIN1:H. While an NMR-based or CryoEM-based physical evidence of direct protein-protein interactions is beyond the scope of present study, our simulation supports the hypothesis that nuclear BIN1:L in OL lineage regulates the cell cycle by inhibiting c-Myc. As the SH3 domain can also bind to tau protein’s proline-rich domain (Sottejeau et al., 2015), we cannot exclude the possibility that BIN1:L in OL lineage may interact with tau proteins and it merits further study (Franzmeier et al., 2019; Ponnusamy et al., 2023; Zhang et al., 2024).

The present study has confirmed that only isoforms 5, 6, 9, 10 and 13 of BIN1:L were detectable in the human brain (Dourlen et al., 2025; Taga et al., 2020). Importantly, BIN1 isoform 9 (ENST00000409400.1 / BIN1-211) is the most abundant transcript in the brain (Dourlen et al., 2025; Taga et al., 2020) and it is highly enriched in the OL lineage (De Rossi et al., 2017; De Rossi et al., 2016). The present study furthers this observation by not only confirming that the transcript of BIN1 isoform 9 is increased in human AD brain, but also showing that the message level of its mouse homologue, Bin1 isoform X20 (ENSMUST00000234857.2 / Bin1-206), is upregulated in the APP/PS1 model. As BIN1 isoform 9 interacts with c-Myc through SH3-MB2 binding, we speculate that increased BIN1 isoform 9 could be mediating nuclear function and regulate OL cell cycle activity in AD.

The present study highlighted the importance of non-neuronal BIN1 that is enriched in myelinating OL. We are the first to describe the nuclear localisation and functions of BIN1 isoforms in OL lineage in human and mouse brain. Nuclear BIN1 expression is associated with cell cycle regulation, and its impairment leads to the perturbation of the c-Myc pathway and contributes to OL degeneration. The loss of OL is a common neuropathology in ageing and AD brain (Bartzokis, 2004; Bartzokis, 2011; Nasrabady et al., 2018; Tse et al., 2018), and such degradation closely tracks with other genetic risk factors like *APOE4* (Cheng et al., 2022). Future investigations are needed to clearly identify the distinct functions and malfunctions of neuron-specific BIN1:H (isoforms 1 - 3) and OL-specific BIN1:L (isoforms 5 - 7, 9, 10, 13) during the pathogenesis of AD and to delineate the effect of *BIN1* genetic variants on neurons and OLs. An important part of this effort will be to delineate the effect of BIN1 genetic AD risk gene variants on neurons and OLs. This expanded view of AD pathogenesis will provide a better mechanistic understanding of exactly how BIN1 variants relate to myelin pathology

## List of Supplementary data

Figure S1 - Homology of human and mouse BIN1 and antibodies-binding sites

Table S1 – Details of postmortem human brain tissues

Table S2 – List of primary antibodies

Table S3 – Primer sequence for gene expression assay

Table S4 – Single cell RNA-seq enquiry

Table S5 – siBin1 RNA-seq output summary

Table S6 – Summary of in silico analysis

