## Supplementary material for "Nuclear BIN1 isoforms regulate c-Myc-mediated cell cycle control in oligodendrocytes": Fig. S1

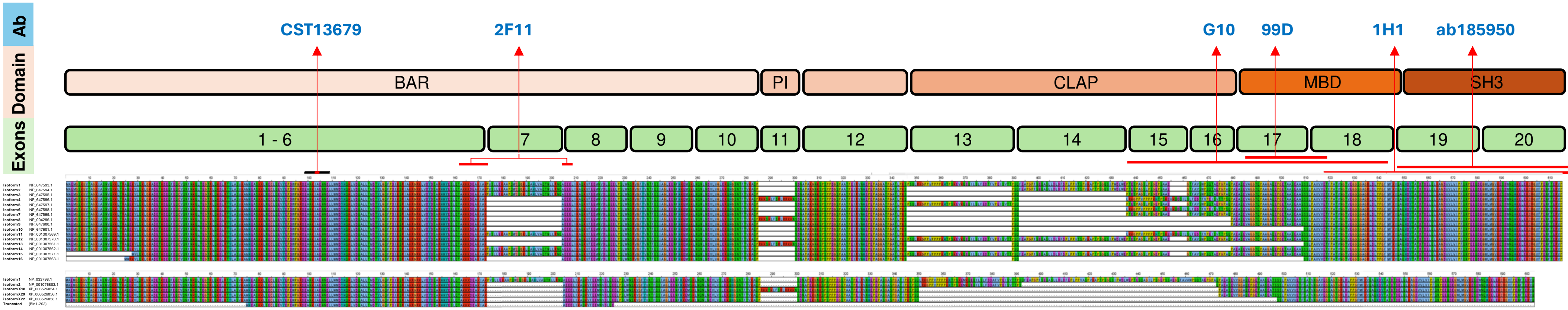


| Species/ clone/name | Cat. no. | Source | Amino acid (UniProt O00499-1) | Amino acid sequence details | Target exon | Human isoform targets | Mouse isoform targets | Isoforms groups | Reference |
|---|---|---|---|---|---|---|---|---|---|
| pAb Rb | CST13679 | Cell signalling | Amino acid residue surrounding alanine 104 | KLNECLQEVYEPDWPGRDEANKIAENNDLLWMDYHQKLVDQALLTMDT YL | Exon 6-10 | All | All | BIN1:H+L | Manufacturer datasheet |
| mAb 2F11 | SC23918 | Santa Cruz | Amino acid 162-173; 205-218 | TAKKKDEAKIAKAEEELIKAQKVFEE | Exon 6-8 junction | 4-10, 13 | 2, X18, X20, X22, Truncated | BIN1:L only | DuHadaway et al 2003 J Cell Biochem |
| mAb 99D | SC13575 | Santa Cruz | Amino acid 323-356 (U68485.1 in year 1997) | SEVAGGTQPAAGAQEPGETAASEAASSSLPAVVV (U68485.1; year 1997) | Exon 17 | 1-9 | 1, 2, X18, X20, X22 | BIN1:H+L | Wechsler-Reya et al 1997 Cancer Res; de Rossi et al 2016 Mol Neurodegen |
| mAb Amphiphysin II (G-10) | sc-74487 | Santa Cruz | Amino acid 421-520 | EPTESPAGSLPSGEPSAAEGTFAVSWPSQTAEPGPAQPAEASEVAGGT QPAAGAQEPGETAASEAASSSLPAVVVETFPATVNGTVEGGSGAGRLDL PPG | Exon 18-19 | 1-3, 5, 7, 13 | Not detectable | BIN1:H+L | Manufacturer datasheet |
| mAb EPR13463-25 | ab185950 | abcam | Amino acid 400 - 593 | PVTSPVKAPTPSGQSIPWDLWEPTESPAGSLPSGEPSAAEGTFAVSWPS QTAEPGPAQPAEASEVAGGTQPAAGAQEPGETAASEAASSSLPAVVVET FPATVNGTVEGGSGAGRLDLPPGFMFKVQAQHDYTATDTDELQLKAGD VVLVIPFQNPEEQDEGWLMGVKESDWNQHKELEKCRGVFPENFTERVP [exact sequence is a commercial proprietary] | Exon 19-20 [estimate] | All | All | BIN1:H+L | Manufacturer communications |
| mAb 1H1 | WH0000274M1 | Millipore/ Sigma | Amino acid 494-593 | VVETFPATVN GTVEGGSGAG RLDLPPGFMFKVQAQHDYTATDTDELQLKA GDVVLVIPFQNPEEQDEGWLMGVKESDWNQHKELEKCRGVFPENFTER VP | Exon 20 | All | All | BIN1:H+L | Manufacturer datasheet |

**Figure S1A - Homology of human and mouse BIN1 and antibodies-binding sites**

```
Score = 29240.0
Length of alignment = 594
Sequence sp|O08539|BIN1_MOUSE/1-588 (Sequence length = 588)
Sequence          NP_647593.1/1-593 (Sequence length = 593)

sp|O08539|BIN1_MOUSE/1-588 MAEMGSKGVTAGKIASNVQKKLTRAQEKVLQKLGKADETKDEQFE
                           |||||||||||||||||||||||||||||||||||||||||||||
         NP_647593.1/1-593 MAEMGSKGVTAGKIASNVQKKLTRAQEKVLQKLGKADETKDEQFE

sp|O08539|BIN1_MOUSE/1-588 QCVQNFNKQLTEGTRLQKDLRTYLASVKAMHEASKKLSECLQEVY
                           |||||||||||||||||||||||||||||||||||||.|||||||
         NP_647593.1/1-593 QCVQNFNKQLTEGTRLQKDLRTYLASVKAMHEASKKLNECLQEVY

sp|O08539|BIN1_MOUSE/1-588 EPEWPGRDEANKIAENNDLLWMDYHQKLVDQALLTMDTYLGQFPD
                           ||.||||||||||||||||||||||||||||||||||||||||||
         NP_647593.1/1-593 EPDWPGRDEANKIAENNDLLWMDYHQKLVDQALLTMDTYLGQFPD

sp|O08539|BIN1_MOUSE/1-588 IKSRIAKRGRKLVDYDSARHHYESLQTAKKKDEAKIAKPVSLLEK
                           |||||||||||||||||||||||||||||||||||||||||||||
         NP_647593.1/1-593 IKSRIAKRGRKLVDYDSARHHYESLQTAKKKDEAKIAKPVSLLEK

sp|O08539|BIN1_MOUSE/1-588 AAPQWCQGKLQAHLVAQTNLLRNQAEEELIKAQKVFEEMNVDLQE
                           |||||||||||||||||||||||||||||||||||||||||||||
         NP_647593.1/1-593 AAPQWCQGKLQAHLVAQTNLLRNQAEEELIKAQKVFEEMNVDLQE

sp|O08539|BIN1_MOUSE/1-588 ELPSLWNSRVGFYVNTFQSIAGLEENFHKEMSKLNQNLNDVLVSL
                           |||||||||||||||||||||||||||||||||||||||||||.|
         NP_647593.1/1-593 ELPSLWNSRVGFYVNTFQSIAGLEENFHKEMSKLNQNLNDVLVGL

sp|O08539|BIN1_MOUSE/1-588 EKQHGSNTFTVKAQPSDNAPEKGNKSPSPPPDGSPAATPEIRVNH
                           |||||||||||||||||||| ||||||| ||||||||||||||||
         NP_647593.1/1-593 EKQHGSNTFTVKAQPSDNAPAKGNKSPS-PPDGSPAATPEIRVNH

sp|O08539|BIN1_MOUSE/1-588 EPEPASGASPGATIPKSPSQLRKGPPVPPPPKHTPSKEMKQEQIL
                           |||||.||.||||.||||||||||||||||||||||||.||||||
         NP_647593.1/1-593 EPEPAGGATPGATLPKSPSQLRKGPPVPPPPKHTPSKEVKQEQIL

sp|O08539|BIN1_MOUSE/1-588 SLFDDAFVPEISVTTPSQFEAPGPFSEQASLLDLDFEPLPPVASP
                           |||.|.||||||||||||||||||||||||||||||.|||||.||
         NP_647593.1/1-593 SLFEDTFVPEISVTTPSQFEAPGPFSEQASLLDLDFDPLPPVTSP

sp|O08539|BIN1_MOUSE/1-588 VKAPTPSGQSIPWDLWEPTESQAGILPSGEPSSAEGSFAVAWPSQ
                           ||||||||||||||||||||| || |||||||.|||.|||.||||
         NP_647593.1/1-593 VKAPTPSGQSIPWDLWEPTESPAGSLPSGEPSAAEGTFAVSWPSQ

sp|O08539|BIN1_MOUSE/1-588 TAEPGPAQPAEASEVVGG------AQEPGETAASEATSSSLPAVV
                           ||||||||||||||| ||      ||||||||||||.||||||||
         NP_647593.1/1-593 TAEPGPAQPAEASEVAGGTQPAAGAQEPGETAASEAASSSLPAVV

sp|O08539|BIN1_MOUSE/1-588 VETFSATVNGAVEGSAGTGRLDLPPGFMFKVQAQHDYTATDTDEL
                           ||||.|||||.|||..|.|||||||||||||||||||||||||||
         NP_647593.1/1-593 VETFPATVNGTVEGGSGAGRLDLPPGFMFKVQAQHDYTATDTDEL

sp|O08539|BIN1_MOUSE/1-588 QLKAGDVVLVIPFQNPEEQDEGWLMGVKESDWNQHKELEKCRGVF
                           |||||||||||||||||||||||||||||||||||||||||||||
         NP_647593.1/1-593 QLKAGDVVLVIPFQNPEEQDEGWLMGVKESDWNQHKELEKCRGVF

sp|O08539|BIN1_MOUSE/1-588 PENFTERVQ
                           ||||||||
         NP_647593.1/1-593 PENFTERVP

Percentage ID = 94.61
```

Yellow = mouse
Green = human

CST13679 Alanine 104; Conserved

2F11 Exon 6-Exon 8 junction; Conserved

G10 Amino acid 421-520; Major shift in mouse (homology = 77.8%)

99D Exon 17; Partially conserved (red font)

ab185950 Amino acid 400-593; Conserved

1H1 Amino acid 494-593; Conserved (underlined)

**Figure S1B - Homology of human and mouse BIN1 and antibodies-binding sites**
