## Supplementary material for "Nuclear BIN1 isoforms regulate c-Myc-mediated cell cycle control in oligodendrocytes": Table S1

| Case | | | | | | Block Details | | AD Neuropathological Changes | |
|---|---|---|---|---|---|---|---|---|---|
| Cohort | # | Gender | Age Range | Condition | PMI (hrs) | Frozen Block | Gross Region | Neocortical amyloid plaques | Neocortical NFTs |
| NIH NeuroBioBank Cohort (Frozen) | NC 1 | F | 70-74 | NC | 19 | A | Frontal Cortex (BA8/9) | Unknown | Unknown |
| | NC 2 | F | 70-74 | NC | 20 | A | Frontal Cortex (BA8/9) | Unknown | Unknown |
| | NC 3 | F | 70-74 | NC | 13 | A | Frontal Cortex (BA8/9) | Unknown | Unknown |
| | NC 4 | M | 70-74 | NC | 12 | A | Frontal Cortex (BA8/9) | Unremarkable | Unremarkable |
| | NC 5 | F | 70-74 | NC | 24 | A | Frontal Cortex (BA8/9) | Unknown | Unknown |
| | NC 6 | M | 70-74 | NC | 15 | A | Frontal Cortex (BA8/9) | Unremarkable | Unremarkable |
| | NC 7 | M | 75-79 | NC | 18 | A | Frontal Cortex (BA8/9) | Mild | Mild |
| | NC 8 | M | 75-79 | NC | 17 | A | Frontal Cortex (BA8/9) | Mild | Mild |
| | DEM 1 | M | 55-59 | Non-AD Dementia | 20.1 | A | Frontal Cortex (BA8/9) | Negative | Negative |
| | DEM 2 | M | 65-69 | Non-AD Dementia | 20 | A | Frontal Cortex (BA8/9) | Remarkable | Remarkable |
| | DEM 3 | F | 55-59 | Non-AD Dementia | 19.7 | A | Frontal Cortex (BA8/9) | Negative | Negative |
| | DEM 4 | F | 80-84 | Non-AD Dementia | 12.8 | A | Frontal Cortex (BA8/9) | Minimal | Negative |
| | DEM 5 | M | 70-74 | Non-AD Dementia | 18.6 | A | Frontal Cortex (BA8/9) | Negative | Negative |
| | DEM 6 | F | 85-90 | Non-AD Dementia | 20 | A | Frontal Cortex (BA8/9) | Negative | Negative |
| | DEM 7 | F | 70-74 | Non-AD Dementia | 26.8 | A | Frontal Cortex (BA8/9) | Negative | Negative |
| | DEM 8 | M | 60-64 | Non-AD Dementia | 19 | A | Frontal Cortex (BA8/9) | Negative | Negative |
| | AD 1 | M | 65-69 | sAD | 5 | A | Frontal Cortex (BA8/9) | Abundant | Abundant |
| | AD 2 | F | 70-74 | sAD | 12 | A | Frontal Cortex (BA8/9) | Abundant | Abundant |
| | AD 3 | F | 75-79 | sAD | 15 | A | Frontal Cortex (BA8/9) | Abundant | Abundant |
| | AD 4 | M | 80-84 | sAD | 9 | A | Frontal Cortex (BA8/9) | Abundant | Abundant |
| | AD 5 | F | 70-74 | sAD | 22 | A | Frontal Cortex (BA8/9) | Abundant | Abundant |
| | AD 6 | F | 75-79 | sAD | 17 | A | Frontal Cortex (BA8/9) | Abundant | Abundant |
| | AD 7 | M | 75-79 | sAD | 3 | A | Frontal Cortex (BA8/9) | Abundant | Abundant |
| | AD 8 | M | 75-79 | sAD | 23 | A | Frontal Cortex (BA8/9) | Abundant | Abundant |

NC: Neurotypical control; DEM: Non-AD Dementia; AD: Sporadic Alzheimer's Disease

**Table S1A - Demographic and histological details of human brain tissue cohorts (NeuroBioBank)**

| Case | | | | | | Block Details | | AD Neuropathological Changes | | |
|---|---|---|---|---|---|---|---|---|---|---|
| **Cohort** | **#** | **Gender** | **Age Range** | **Condition** | **PMI (hrs)** | **FFPE Block** | **Gross Region** | **NFT (Braak)** | **Neuritic Plaque CERAD** | **NIA-RI AD Likelihood** |
| **University of Pittsburgh UPMC ADRC cohort (FFPE)** | NC 1 | M | 80-84 | NC | 5.0 | A | Frontal Cortex (BA8/9) | I | N.A. | Low |
| | NC 2 | M | 70-74 | NC | 3.0 | A | Frontal Cortex (BA8/9) | II | N.A. | N.A. |
| | NC 3 | M | 85-89 | NC | 7.0 | A | Frontal Cortex (BA8/9) | II | Frequent | Low |
| | NC 4 | M | 60-64 | NC | 6.5 | A | Frontal Cortex (BA8/9) | 0 | N.D. | Low |
| | NC 5 | M | 75-79 | NC | 6.0 | A | Frontal Cortex (BA8/9) | II | N.D. | Low |
| | NC 6 | F | 75-79 | NC | 10.5 | A | Frontal Cortex (BA8/9) | II | N.D. | Low |
| | NC 7 | M | 70-74 | NC | 4.5 | A | Frontal Cortex (BA8/9) | I | Sparse | Low |
| | NC 8 | M | 90-94 | NC | 4.0 | A | Frontal Cortex (BA8/9) | II | N.D. | Low |
| | NC 9 | M | 75-79 | NC | 14.0 | A | Frontal Cortex (BA8/9) | II | Sparse | Low |
| | AD 1 | M | 75-79 | sAD | 2.0 | A | Frontal Cortex (BA8/9) | IV | Frequent | Intermediate |
| | AD 2 | M | 75-79 | sAD | 15.0 | A | Frontal Cortex (BA8/9) | IV | Frequent | Intermediate |
| | AD 3 | M | 85-89 | sAD | 4.0 | A | Frontal Cortex (BA8/9) | IV | Frequent | Intermediate |
| | AD 4 | M | 80-84 | sAD | 12.0 | A | Frontal Cortex (BA8/9) | IV | Frequent | Intermediate |
| | AD 5 | M | 80-84 | sAD | 6.0 | A | Frontal Cortex (BA8/9) | IV | Frequent | Intermediate |
| | AD 6 | M | 85-89 | sAD | 4.0 | A | Frontal Cortex (BA8/9) | IV | Frequent | Intermediate |
| | AD 7 | M | 90-94 | sAD | 6.0 | A | Frontal Cortex (BA8/9) | III | Frequent | Intermediate |
| | AD 8 | F | 85-89 | sAD | 6.0 | A | Frontal Cortex (BA8/9) | IV | Frequent | Intermediate |
| | AD 9 | M | 85-89 | sAD | 9.0 | A | Frontal Cortex (BA8/9) | III | Frequent | Intermediate |
| | AD 10 | M | 85-89 | sAD | 3.0 | A | Frontal Cortex (BA8/9) | VI | Frequent | High |
| | AD 11 | M | 75-79 | sAD | 4.0 | A | Frontal Cortex (BA8/9) | VI | Frequent | High |
| | AD 12 | F | 75-79 | sAD | 4.0 | A | Frontal Cortex (BA8/9) | VI | Frequent | High |
| | AD 13 | M | 80-84 | sAD | 7.5 | A | Frontal Cortex (BA8/9) | VI | Frequent | High |
| | AD 14 | M | 85-89 | sAD | 7.0 | A | Frontal Cortex (BA8/9) | V | Frequent | High |
| | AD 15 | F | 75-79 | sAD | 2.0 | A | Frontal Cortex (BA8/9) | VI | Frequent | High |
| | AD 16 | M | 80-84 | sAD | 4.0 | A | Frontal Cortex (BA8/9) | VI | Frequent | High |
| | AD 17 | F | 80-84 | sAD | 2.0 | A | Frontal Cortex (BA8/9) | VI | Frequent | High |
| | AD 18 | F | 75-79 | sAD | 3.0 | A | Frontal Cortex (BA8/9) | VI | Frequent | High |
| | AD 21 | F | 55-59 | fAD | 9.0 | A | Frontal Cortex (BA8/9) | VI | Definite | High |
| | AD 22 | F | 45-49 | fAD | 18.0 | A | Frontal Cortex (BA8/9) | VI | Definite | High |
| | AD 23 | F | 45-49 | fAD | 6.0 | A | Frontal Cortex (BA8/9) | VI | Definite | High |
| | AD 24 | M | 50-54 | fAD | 3.0 | A | Frontal Cortex (BA8/9) | VI | Definite | High |
| | AD 25 | M | 50-54 | fAD | N.A. | A | Frontal Cortex (BA8/9) | VI | Definite | High |
| | AD 26 | F | 45-49 | fAD | 10.0 | A | Frontal Cortex (BA8/9) | VI | Definite | High |
| | AD 27 | F | 50-54 | fAD | 4.0 | A | Frontal Cortex (BA8/9) | VI | Definite | High |

NC: Neurotypical control; AD: Alzheimer's Disease; sAD: sporadic Alzheimer's Disease; fAD: familial Alzheimer's Disease

**Table S1B - Demographic and histological details of human brain tissue cohorts (Pittsburgh)**
