## Supplementary material for "Nuclear BIN1 isoforms regulate c-Myc-mediated cell cycle control in oligodendrocytes": Table S2

| Antibody | Host species | Catalog no. | Lot no. | Source | WB Dilution | IHC Dilution | ICC Dilution |
|---|---|---|---|---|---|---|---|
| Anti-MAG Antibody, clone 513, Alexa Fluor 488 Conjugate | mouse | MAB1567A4 | 4079388 | Merck Millipore | 1:1000 | N/A | 1:500 |
| Anti-NeuN Antibody, clone A60, Cy3 Conjugate | mouse | MAB377C3 | 2943768 | Merck Millipore | 1:1000 | 1:500 | N/A |
| Anti-Olig2 Antibody, clone 211F1.1, Alexa Fluor 488 Conjugate | mouse | MABN50A4 | 3756534 / 4067709 / 4057431 / 3822360 | Sigma | N/A | N/A | 1:500 |
| APC, clone: CC1 | mouse | MABC200 | 3698630 | Merck Millipore | N/A | 1:500 | N/A |
| BIN1 | rabbit | 13679S | 1 | Cell signaling Technology | 1:500 | N/A | N/A |
| BIN1 (3B6A4) | mouse | MA5-31651 | YA3815225 | Invitrogen | 1:500 | N/A | N/A |
| BIN1 (EPR13463-25) | rabbit | ab185950 | GR3258620-1 / 1006598-2 / 1006598-9 | Abcam | 1:500 | 1:250 | 1:250 |
| BIN1 / Amphiphysin II (2F11) | mouse | sc-23918 | H2823 / G1921 | Santa Cruz | 1:500 | 1:250 | 1:250 |
| BIN1 / Amphiphysin II (99D) | mouse | sc-13575 | D2723 | Santa Cruz | 1:500 | 1:250 | 1:250 |
| BIN1 / Amphiphysin II (G10) | mouse | sc-74487 | J0509 | Santa Cruz | 1:500 | 1:250 | 1:250 |
| BIN1, clone 1H1 | mouse | WH0000274M1-100UG | GC191-1H21 | Sigma | 1:500 | N/A | N/A |
| c-Myc (9E10) | mouse | MA1-980 | XD347120 | Invitrogen | N/A | N/A | 1:250 |
| c-Myc (D84C12) | rabbit | 5605S | 12, 16 | Cell signaling Technology | 1:1000 | N/A | N/A |
| CSPG4 (NG2) | rabbit | AB5320 | 3118137 / 3457641 / 2887981 | Merck Millipore | N/A | N/A | 1:400 |
| Cyclin D1 (72-13G) | mouse | sc-450 | F2719 | Santa Cruz | N/A | N/A | 1:250 |
| GAPDH (6C5) | mouse | ab8245 | GR3438148-1 / GR3275542-6 / 1035914-6 | Abcam | 1:10000 | N/A | N/A |
| MAG (D4G3) XP (R) | rabbit | 9043S | 2 | Cell signaling Technology | N/A | N/A | 1:250 |
| Olig2, clone 211F1.1 | mouse | MABN50 | 3756534 | Merck Millipore | N/A | N/A | 1:250 |
| Purified anti-β-Amyloid, 1-16 Antibody (6E10) | mouse | 803001 | B378932 | BioLegend | N/A | 1:500 | N/A |

WB: Western Blotting; IHC: Immunofluorescence; ICC: Immunocytochemistry

**Table S2. List of primary antibodies and dilution used in this study**
