## Supplementary material for "Nuclear BIN1 isoforms regulate c-Myc-mediated cell cycle control in oligodendrocytes": Table S3

| NCBI Isoform # | Ensembl Protein Name | ENST (Ensembl spliced transcripts ) | Design | Forward primer | Forward primer - target exonic region | Reverse primer | Reverse primer - target exonic region |
|---|---|---|---|---|---|---|---|
| **HUMAN** | | | | | | | |
| **1** | BIN1-202 | ENST00000316724 | This study | CCAAGTCCCCATCTCAGCTC | on exon 12 | CGGGGCCTCAAACTGGG | span the border of exon13 & exon14 |
| **2** | BIN1-207 | ENST00000357970 | This study | CCCAAGTCCCCATCTCAGTTT | span the border of exon12 & exon14 | CCCAGAGGTCCCATGGAATTG | on exon 15 |
| **3** | BIN1-205 | ENST00000351659 | This study | CAAGTCCCCATCTCAGTCAA | span the border of exon12 & exon15 | CACGACAGCAGGAAGAGAG | on exon 18 |
| **4** | BIN1-201 | ENST00000259238 | This study | GCGCAGAAAGAAGAACAGTG | span the border of exon11 & exon12 | TCTGCTGGCTGGGAGG | span the border of exon13 & exon17 |
| **5** | BIN1-203 | ENST00000346226 | This study | GAAGTCAAGCAGGAGCAGAT | on exon 13 | CTCTCTGTGGGCTGGGA | span the border of exon13 & exon16 |
| **4&5&6** | BIN1-201,-203,-209 | ENST00000259238, ENST00000346226, ENST00000393040 | This study | ACCCCCTCCCAGCCA | span the border of exon13 & exon17 | TTGAACATGAAACCTGGGGG | on exon 18 |
| **7** | BIN1-210 | ENST00000393041 | This study | CCAAAATTGCCAAGGCCG | span the border of exon6 & exon8 | GACTCTCTGTGGGCTGAG | span the border of exon12 & exon16 |
| **8&9** | BIN1-206,-211 | ENST00000352848, ENST00000409400 | This study | CAAGTCCCCATCTCAGCCAG | span the border of exon12 & exon17 | TGAACATGAAACCTGGGGG | on exon 18 |
| **10** | BIN1-204 | ENST00000348750 | This study | CCCAGCCCAGTGACAAC | span the border of exon10 & exon12 | AGGAAGAGAGCTCTGAGATG | span the border of exon12 & exon18 |
| **13** | BIN1-208 | ENST00000376113 | This study | AAGAAGAACAGTGACAACGC | span the border of exon11 & exon12 | GGAAGAGAGCTCTGAGATGG | span the border of exon12 & exon18 |
| **1-3** | BIN1-202,-207,-205 | BIN1: L / D7 | de Ross et al 2016 Mol. Neurodegen | CCTGCTGTGGATGGATTACC | at the start of exon 5 | TCCTCCTCGGCCTTGGC | span the board of exon 6 and exon 8, skipping exon 7 |
| **4-10, 13** | BIN1-201 to -204 & -208 | BIN1: L / Ex7 | de Ross et al 2016 Mol. Neurodegen | CCTGCTGTGGATGGATTACC | at the start of exon 5 | GCTTTCTCAAGCAGCGAGAC | on exon 7 |
| **MOUSE** | | | | | | | |
| **All** | | All | This study | GATGCACGAAGCCTCCAAGA | Primer pairs within exon 4-6 | GGAACTGGCCTAGGTAGGTG | Primer pairs within exon 4-6 |
| **1** | Bin1-201 | ENSMUST00000025239.9 | This study | CACACCCTCTGGTCAGTCAA | On the boundary on exon 13 | AGAGCTGGAGGTTGCTTCAC | On the boundary on exon 14 |
| **2** | Bin1-202 | ENSMUST00000091967.13 | This study | AGAAAGGGAACAAGAGCCCG | On the boundary on exon 13 | TCTGTGGGCTGGGAGG | On the boundary on exon 17 |
| **X20** | Bin1-206 | ENSMUST00000234857.2 | This study | CAAGTCCCCATCTCAGCCAG | On the boundary on exon 12 | GCTGGGCTTGAACCTTGAAC | On the boundary on exon 17 |
| **X22** | Bin1-205 | ENSMUST00000234496.2 | This study | GGGAGCAACACCTTCACAGT | On the boundary on exon 12 | CCGGAAGAGAGCTCTGAGATG | On the boundary on exon 18 |

**Table S3. Human and mouse BIN1 primers design**
