## Supplementary material for "Nuclear BIN1 isoforms regulate c-Myc-mediated cell cycle control in oligodendrocytes": Table S6

| | | Ensembl Protein Name | NCBI Isoform # | NCBI Ref ID | UniProt Ref ID | Length (a.a.) | Biotype | AlphaFold Model ID | c-Myc Docking results | Hydrophobic-favoured Centre Energy (KJ mol$^{-1}$) |
|---|---|---|---|---|---|---|---|---|---|---|
| **BIN1** | | | | | | | | | | |
| **Human** | **BIN1:H** | BIN1-202 | Isoform 1 | NP_647593.1 | O00499-1 | 593 | Protein coding | AF-O00499-F1-model_v4.pdb | OKAY | -2464 |
| | | BIN1-207 | isoform 2 | NP_647594.1 | O00499-5 | 550 | Protein coding | AF-A0A024RAF6-F1-model_v4.pdb | OKAY | -1735 |
| | | BIN1-205 | isoform 3 | NP_647595.1 | O00499-3 | 506 | Protein coding | AF-A0A024RAJ3-F1-model_v4.pdb | OKAY | -1806 |
| | **BIN1:L** | BIN1-201 | isoform 4 | NP_647596.1 | O00499-11 | 497 | Protein coding | AF-A0A024RAJ2-F1-model_v4.pdb | Fail | / |
| | | BIN1-203 | isoform 5 | NP_647597.1 | O00499-2 | 518 | Protein coding | AF-A0A024RAF0-F1-model_v4.pdb | Fail | / |
| | | BIN1-209 | isoform 6 | NP_647598.1 | O00499-6 | 482 | Protein coding | AF-A0A024RAI5-F1-model_v4.pdb | Fail | / |
| | | BIN1-210 | isoform 7 | NP_647599.1 | O00499-4 | 475 | Protein coding | AF-A0A024RAI4-F1-model_v4.pdb | Fail | / |
| | | BIN1-206 | isoform 8 | NP_004296.1 | O00499-8 | 454 | Protein coding | AF-A0A024RAE9-F1-model_v4.pdb | Fail | / |
| | | BIN1-211 | isoform 9 | NP_647600.1 | O00499-7 | 439 | Protein coding | AF-A0A024RAG8-F1-model_v4.pdb | OKAY | -2163 |
| | | BIN1-204 | isoform 10 | NP_647601.1 | O00499-9 | 409 | Protein coding | AF-A0A024RAI6-F1-model_v4.pdb | OKAY | -1922 |
| | | BIN1-208 | isoform 13 | NP_001307561.1 | O00499-10 | 424 | Protein coding | AF-A0A024RAG9-F1-model_v4.pdb | OKAY | -1618 |
| **Mouse** | **Bin1:H** | Bin1-201 | Isoform 1 | NP_033798.1 | O08539-1 | 588 | Protein coding | AF-O08539-F1-model_v4.pdb | OKAY | -2142 |
| | **Bin1:L** | Bin1-202 | isoform 2 | NP_001076803.1 | Q6P1B9 | 477 | Protein coding | AF-Q6P1B9-F1-model_v4.pdb | OKAY | -1637 |
| | | Bin1-205 | isoform X22 | XP_006526058.1 | A0A3Q4EBK4 | 410 | Protein coding | AF-A0A3Q4EBK4-F1-model_v4.pdb | Fail | / |
| | | Bin1-203 | N/A | N/A | A0A3Q4EBR8 | 120 | Protein coding (Truncated) | AF-A0A3Q4EBR8-F1-model_v4.pdb | OKAY | -1784 |
| **c-Myc** | | | | | | | | | | |
| **Human** | **C-MYC** | MYC-202 | C-MYC | N/A | P01106 | 439 | Protein coding | AF-P01106-F1-model_v4.pdb | / | / |
| **Mouse** | **c-myc** | Myc-205 | c-myc | N/A | P01108 | 439 | Protein coding | AF-P01108-F1-model_v4.pdb | / | / |

**Table S6 – Summary of in silico analysis**
